# Social microbiome transmission predicts microbial specialization and host lifespan in a wild primate

**DOI:** 10.64898/2026.04.29.721577

**Authors:** Reena Debray, Caroline C. Dickson, Shasta E. Webb, Pamela Ferretti, Alex Meloimet, Jack Gilbert, Susan C. Alberts, Ran Blekhman, Elizabeth A. Archie, Jenny Tung

## Abstract

Social interactions are proposed to provide reliable routes for microbial transmission between animals, facilitating animal-microbiome co-evolution. However, microbiome transmission remains challenging to measure in wild populations. Here we combine behavioral observations of wild baboons with repeated strain-resolved metagenomic profiling to identify individual gut microbial species that follow a dominant mode of social transmission. In an 18-year metagenomic time series from the same population, baboons with higher levels of socially transmitted species lived longer than those with lower levels of socially transmitted species. Socially transmitted species were also more stable and persistent within baboons, yet had narrower host ranges outside of baboons. Thus, social transmission is not only detectable in free-living primates, but may play a special role in both host and microbial fitness.

## INTRODUCTION

All animals live in symbiotic relationships with microorganisms, which they acquire from either the surrounding environment or other individuals. Social interactions can thus shape microbiome transmission, with important implications for both hosts and their microbes (*1*–*3*). Microorganisms that pass directly between hosts through social transmission likely experience different selection regimes than those that cycle through non-host environments (*4*). For hosts, social behaviors that facilitate transmission may promote microbiome stability, with consequences for individual health (*5*) and long-term host-microbe co-diversification (*6, 7*).

Experimental studies support the importance of social transmission under laboratory conditions (*8*–*10*). In zebrafish, artificial selection for host-to-host transmission selects for bacteria that survive better within hosts (*11*), while in bumble bees, a socially transmitted bacterium improves host resistance to a protozoan parasite challenge (*12*). Outside of the lab, however, the evolutionary consequences of social transmission remain largely unknown. In part, this is because evidence for social transmission relies on species- or strain-sharing patterns that could be driven by similar lifestyles, ages, or diets among social partners rather than transmission *per se* (*13*–*15*). Consequently, whether socially transmitted microbes are well-adapted to their hosts, or whether their presence predicts host fitness, have not been testable.

To do so here, we used a novel study design that pairs dense, longitudinal gut microbiome sampling with concurrent observations of life history and behavior. We focused on a population of wild baboons in the Amboseli ecosystem of southern Kenya, which has been studied continuously since 1971 (*16*) and where previous work has identified elevated gut microbiome similarity among social partners (*3, 17*). We began by conducting a concentrated 3-month sampling effort to track microbial strains moving between adult baboons. This approach allowed us to resolve both individual transmission events and pinpoint specific species where transmission closely follows a grooming-based social network. Next, using an 18-year longitudinal metagenomic dataset from the same population, we tested whether social transmission predicts fitness proxies for either microbes or their hosts. Finally, to investigate the consequences of social transmission for long-term host-microbe evolution, we evaluated whether social transmission explains co-diversification patterns across primate lineages.

## RESULTS

### High-resolution tracking of microbial transmission in a natural primate population

To identify socially transmitted gut microbiome strains, we performed strain-resolved metagenomic analysis of repeated samples from all adult baboons in the Amboseli study population. Over a period of 79 days, we collected 410 fecal samples from 132 adult baboons living in 5 social groups (median=3 samples/individual; Figure **1a**). We generated shotgun metagenomes and competitively mapped the resulting reads to a custom database of 4,712 species-representative bacterial and archaeal genomes (median 36.0 million reads/sample, Figure **1b**, Table **S1**,**S2**). For each individual, a median of 145 species mapped to this database, of which 76.5 species met our requirements for subsequent strain-level profiling (≥50% of the genome represented at ≥5x coverage). Consistent with previous findings (*17*–*19*), microbiome composition was more similar among baboons from the same social group than baboons from different groups (PERMANOVA, F=1.143, p=0.001; Figure **S1**).

**Figure 1.**
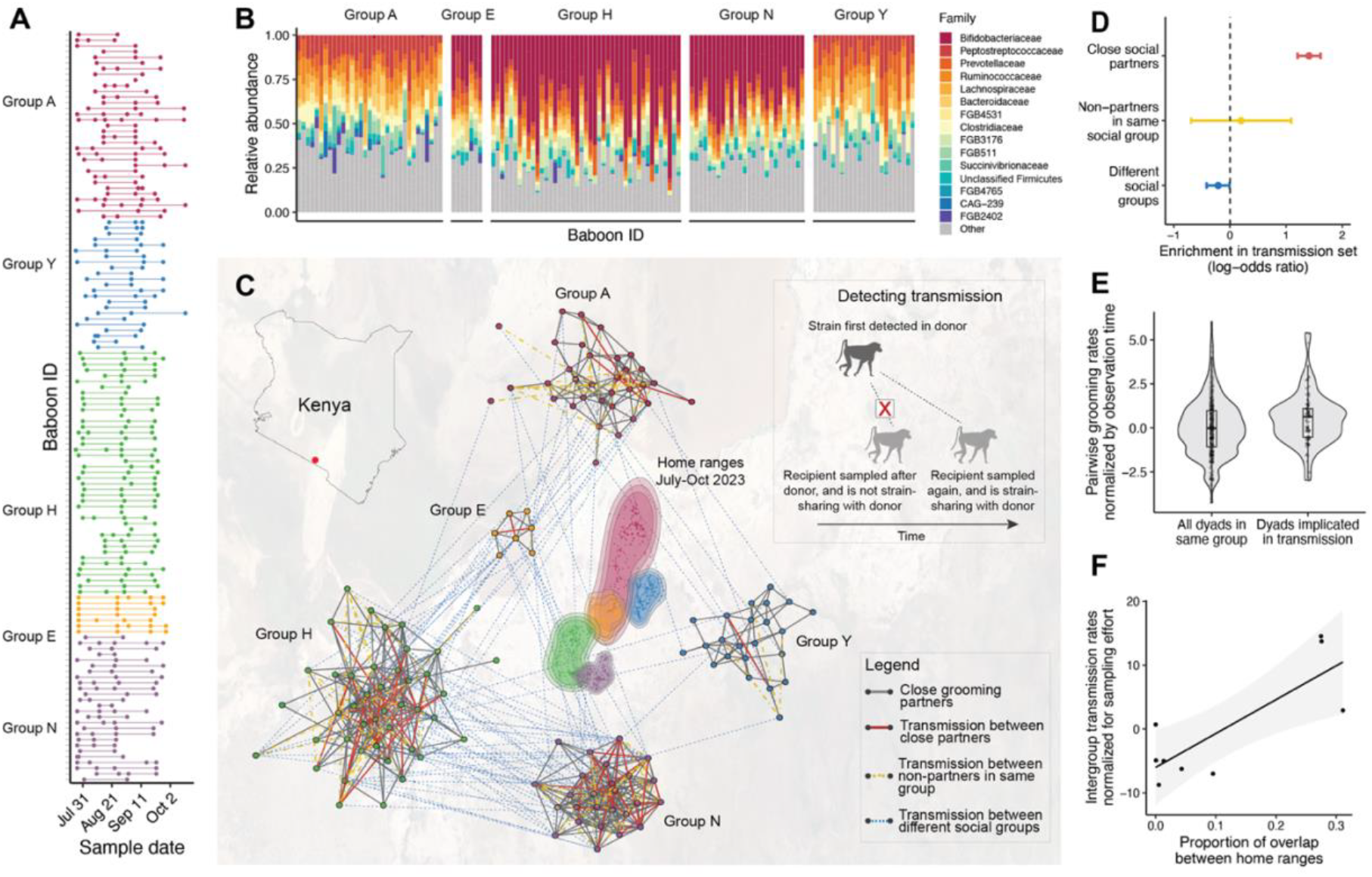
Strain tracking in metagenomes reveals transmission dynamics within baboon social groups. **(a)** Fecal sample collection in the 3-month dataset. Each horizontal line represents an individual baboon; colors represent different social groups. **(b)** Relative abundances of the top 15 microbial families, calculated as the proportion of genomic coverage attributed to each family. **(c)** Home ranges of the five baboon social groups (center) in this study, constructed as autocorrelated kernel density estimates based on regular GPS recordings of group movements. The contours of each home range represent the lower bound, the estimate, and the upper bound of the 95% contour, respectively. Networks represent grooming-based social networks; colored lines indicate microbial transmission events. Inset: to identify a transmission event, we required that strain-sharing involved (i) a sharing event between a unique potential donor and a potential recipient collected after the sample from the donor, and (ii) one or more “intermediate” samples from the recipient collected before strain-sharing was identified but after the donor sample. **(d)** Strain transmission events were enriched between close grooming partners (top 20%) compared to other members of the same group or members of different groups (Fisher’s exact test, log_2_(OR)=1.41, p<0.001). **(e)** Within groups, dyads implicated in transmission spent more time grooming relative to all within-group dyads (logistic regression, ß=0.121, p=0.017). **(f)** Intergroup transmission events occurred more frequently between groups with greater home range overlap (dyadic regression, ß=4.19, p<0.001).

To track strain transmission, we first identified inter-individual strain sharing, in which two baboons carried near-identical populations of the same microbial species. We measured genomic similarity using both a nucleotide identity-based method (*20*) and a genome synteny-based method (*21*). The two methods returned similar strain sharing rates per sample pair, but implicated different microbial species, reflecting their sensitivity to different modes of microbial evolution (Figure **S2**). Combined, we detected 20,061 strain sharing events in the three-month period. We identified new strain acquisitions when: (i) strain sharing occurred between one individual (the donor) and a later-in-time sample from another individual (the recipient), and (ii) one or more additional “intermediate” samples was available from the recipient *before* the sample with the shared strain was collected for the recipient, but *after* the sample with the shared strain was collected for the donor. These criteria indicate a new strain acquisition in the recipient (Figure **1c** **inset, S3**). We focused on transmission events with only one possible donor to confidently assign donor-recipient pairs and directionality.

In total, we observed 206 new strain acquisition events involving 108 individuals and 214 unique dyads (Table **S3**). Detection of transmission was not related to sequencing depth (binomial regression, p=0.544, Methods, Figure **S4, S5**). Immigrant males (i.e., those living in a different social group than they were born in; in the Amboseli baboons, males are the dispersing sex) were enriched for the transmission of new strains, suggesting that changes in social context present opportunities for novel strain acquisition (Fisher’s exact test, log_2_(OR)=0.627, p<0.001; Figure **S6, S7**).

To resolve transmission pathways, we next identified social partners based on the top 20% of grooming frequencies during July-December 2023, which brackets the fecal sampling period. Grooming is the most common form of physical contact in baboons and many other social primates, making it a likely mechanism for microbial transmission. Transmission events were highly enriched between close grooming partners relative to their frequency in the population (Fisher’s exact test, log_2_(OR)=1.41, p<0.001; Figure **1d, 1e**), but not between group-mates who were not grooming partners (log_2_(OR)=0.123, p=0.634). Transmission was depleted between members of different social groups (log_2_(OR) = -0.211, p=0.044; Figure **1d**). Baboons do not typically engage in affiliative social interactions across groups (*22*), and no males dispersed between groups during our study period, suggesting that cases of between-group transmission arise from environmental intermediates (e.g., water holes or sleeping groves). Consistent with this hypothesis, intergroup transmission was elevated between groups with overlapping home ranges and groups that used the same sleeping groves (dyadic regression on Bhattacharyya overlap in home range distributions and transmission rate, estimate=4.19, p<0.001; Figure **1f**; for sleeping groves: estimate=0.77, p<0.001; Figure **S8**).

### Socially transmitted species show hallmarks of host specialization

Some host-associated microbes are generalists that can live in environmental reservoirs between hosts, while others are host specialists (*6, 11, 23*). We tested whether social transmission, which implies continuous residence in hosts, selects for host specialization. To do so, we assigned a dominant transmission mode (≥3 transmission events, with the majority following the same mode) to the 42 microbial species that were either (i) consistently transmitted between close social partners (socially transmitted, ST; n=8 species) or (ii) consistently transmitted between non-social partners (either within groups, NST_within_ [n=9 species], or between groups, NST_between_ [n=25 species]) (Figure **2a**).

**Figure 2.**
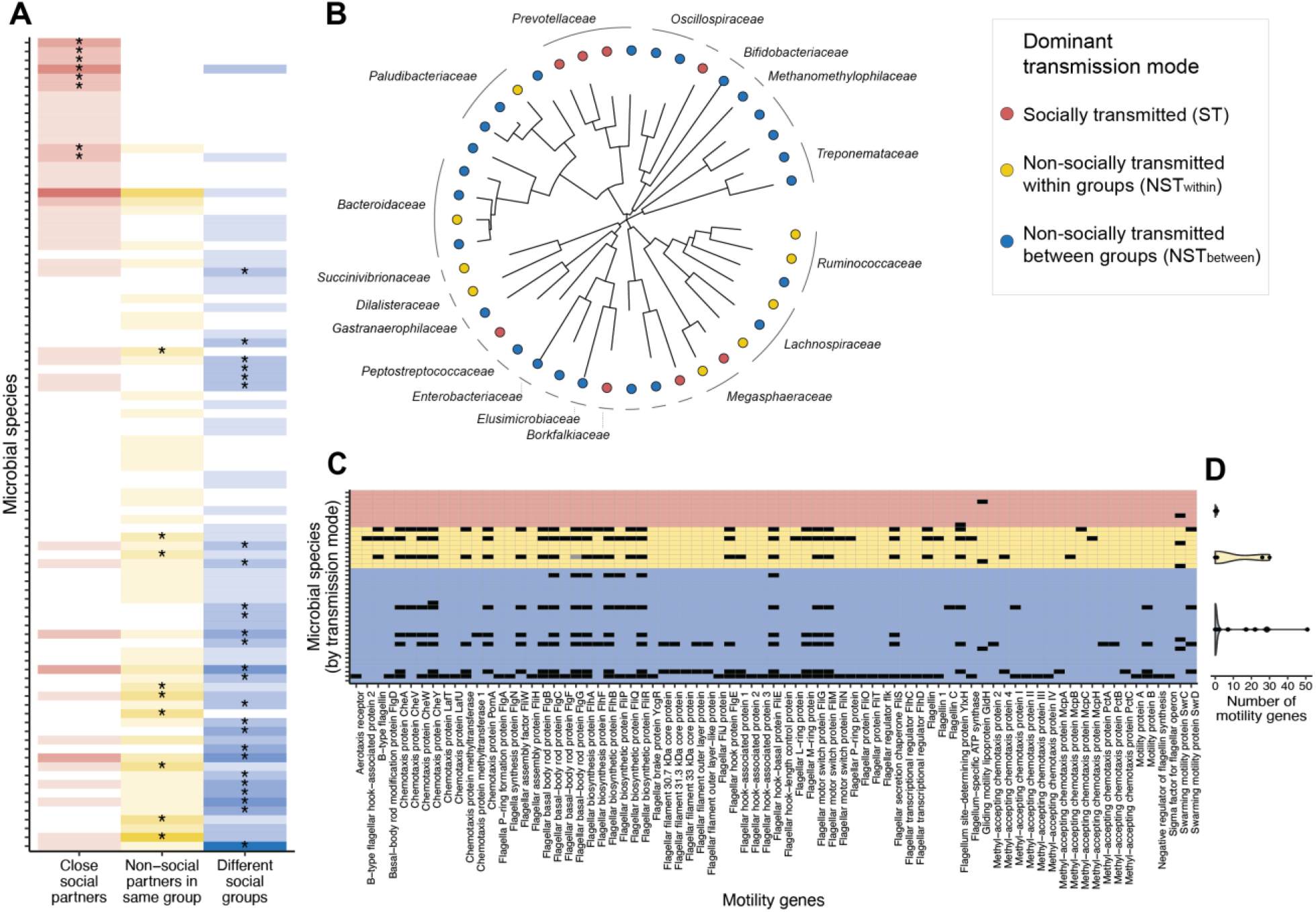
Traits and genome content of socially and non-socially transmitted species. **(a)** Microbial species detected in transmission (n=206 events transmission events involving 92 species) and annotated according to whether the transmission event occurred between close social partners (red), between non-close social partners in the same social group (yellow), or between different social groups (blue). Intensity of shading indicates the number of transmission events (at most six in any species-category combination). Asterisks indicate the 42 species that were implicated in transmission in at least three instances with a clear majority mode (i.e., no ties). **(b)** Phylogenetic tree of 42 microbial species with dominant transmission modes indicated by tip color. Family-level taxonomic classifications are indicated where available. **(c)** Presence or absence of genes involved in motility (chemotaxis, aerotaxis, and flagellar traits) for the 42 species shown in panel B. Black squares indicate an intact gene, while grey squares indicate a gene flagged as a likely pseudogene. **(d)** Total numbers of genes involved in motility by microbial species and dominant transmission mode.

These 42 species were phylogenetically diverse, although social transmission was particularly concentrated within the genus *Prevotella* (3 of 8 ST species; Figure **2b**; Table **S4**). We observed an asymmetric trade-off between transmission modes. Species transmitted more often between social partners were less likely to ever be transmitted between non-social partners (binomial model, coefficient=-0.745, p=0.0069), whereas the frequency of transmission between non-social partners was unrelated to whether microbes were implicated in social transmission (coefficient=0.088, p=0.504, Figure **S9**). This finding implies two distinct microbial strategies: specialists that rely extensively on social transmission, and generalists that can disperse through social or non-social routes.

To further test for host specialization in socially transmitted species, we compared the genomes of ST and NST species. ST and NST species did not differ in genome size, gene pseudogenization rates, predicted oxygen tolerance, or predicted temperature preference (all p>0.50; Figure **S10, S11**). However, the genomes of socially transmitted species were strongly depleted of genes involved in motility (t-test, t=3.03, p=0.005, Figure **2c, 2d**). ST species also carried marginally fewer genes involved in sporulation and toxin-antitoxin systems, both thought to aid in microbial survival outside of hosts (*24, 25*) (sporulation t=1.80, p=0.086; toxin-antitoxin t=1.92, p=0.063; Figure **S10**).

### Socially transmitted species persist longer in hosts than non-socially transmitted species

For host-associated microbes, high fitness within hosts is linked to stable abundance and persistence (*26*). We hypothesized that ST species would exhibit higher within-host fitness as a result of continuous residence in hosts. To test this hypothesis, we turned to an independent, 18-year metagenomic dataset from the same baboon population, representing 5,910 fecal samples collected from 474 individuals between 2000-2018 (median=10 samples/individual, ∼5.5 months between samples: Figure **3a**, **S12**). Metagenomes were sequenced to a median depth of 51.2 million reads/sample (Table **S5**). All 42 ST, NST_within_, and NST_between_ species annotated in the 3-month dataset were also detected in the 18-year dataset.

**Figure 3.**
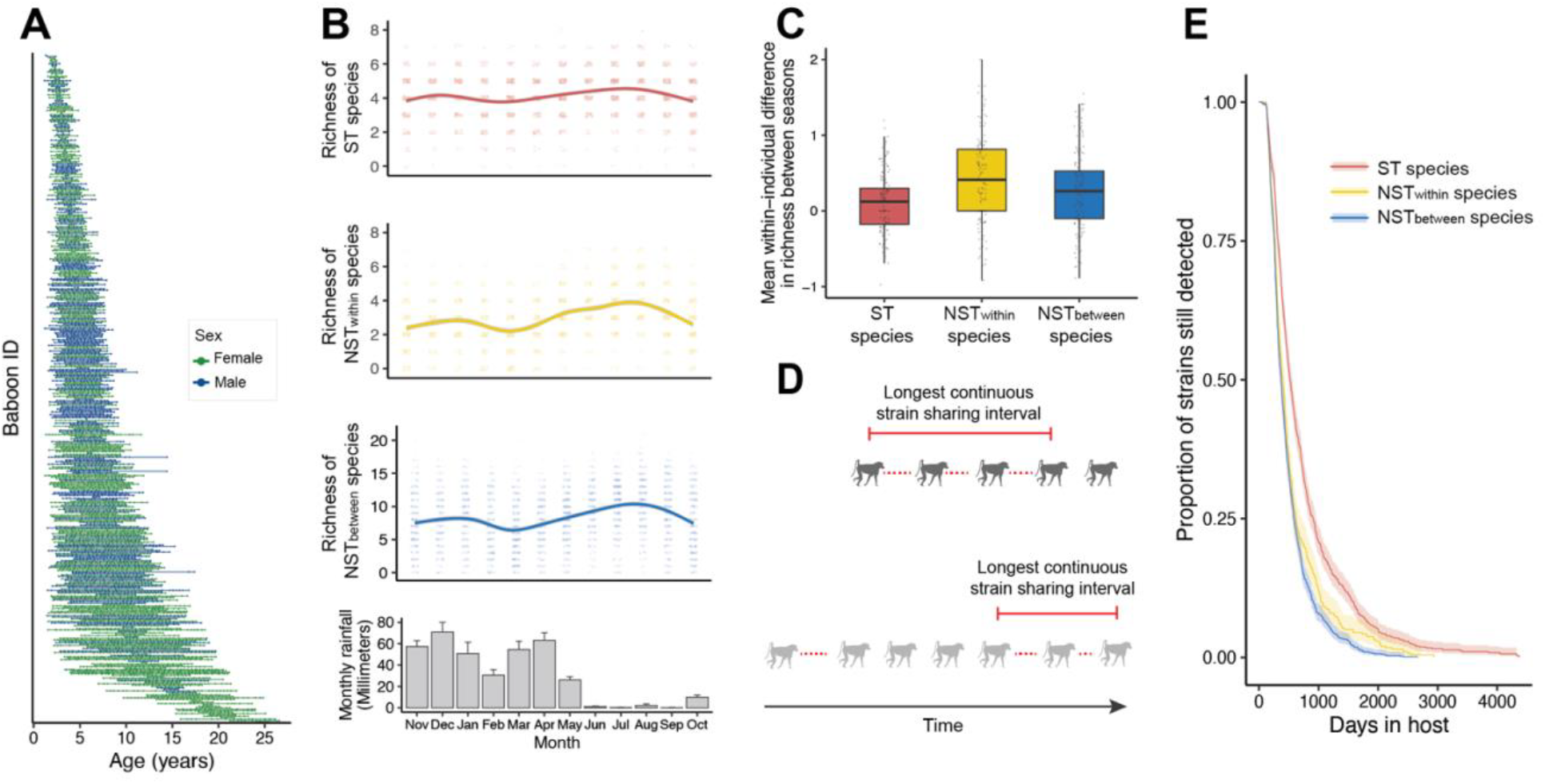
Distribution and ecology of socially and non-socially transmitted species. **(a)** Schedule of fecal sample collection for the 18-year shotgun longitudinal dataset. Each horizontal line represents an individual baboon and colors represent sexes. **(b)** Richness of ST, NST_within_, or NST_between_ species per sample, aligned with monthly mean rainfall in the Amboseli region. NST species varied cyclically and tended to increase following the two annual wet seasons. **(c)** Within-individual differences between average species richness in the wet and dry seasons varied more for NST species than for ST species (ST species versus NST_within_: t=-4.754, p<0.001; ST versus NST_between_: t=-3.333, p=0.001). **(d)** Microbial persistence was calculated for each species-individual combination as the maximum interval for which a strain was continuously shared between serial samples collected from the individual. **(e)** ST species had longer persistence durations than NST species (hazard ratio for drop-outs of ST species = 0.548 compared to NST species, p=0.038).

Despite little change in the abundances and prevalences of ST, NST_within_, or NST_between_ species across years, many of these species exhibited cyclical dynamics within years, coinciding with seasonal changes in the ecosystem (Figure **3b**, Figure **S13, S14**). However, within individuals, ST species richness was more stable than NST_within_ and NST_between_ species (paired t-test ST species versus NST_within_: t=-4.754, p<0.001; ST versus NST_between_: t=-3.333, p=0.001; Figure **3c**). The larger fluctuations of NST species with changes in temperature and rainfall support the interpretation that these species are commonly acquired via environmental intermediates.

To test whether socially transmitted microbial species persist longer within hosts, we defined persistence as the maximum interval over which all consecutive samples from an individual baboon shared strains with each other (Figure **3d**). Most new strains did not persist beyond 6-8 months, although a minority (14.7%) of microbial species had average within-host strain persistences ≥1 year. Per-species persistence values were correlated across hosts, suggesting that they reflect a consistent measure of within-host microbial fitness (Figure **S15**). On average, ST species persisted within individuals for 522 days after first detection, while NST_within_ and NST_between_ species persisted for a median of 399 days and 371 days, respectively (Cox model hazard ratio [HR] for drop-outs of ST species=0.548 compared to NST species, p=0.038; Figure **3e**). In addition, ST species, but not NST species, were enriched among the 51 “long-persisters” with median durations exceeding 1 year (Fisher’s exact test, ST log_2_(OR) = 1.818, p=0.019; NST_within_ log_2_(OR) = 0.520, p=0.623; NST_between_ log_2_(OR) ratio = 0.666, p=0.224; Table **S7**). We obtained qualitatively unchanged results when focusing on a small dataset of very densely sampled baboons (9 individuals sampled every 2-4 weeks for 13 years), which is less vulnerable to drop-out and recolonization events than our primary data set (Figure **S16**).

Finally, because overall strain sharing rates are often used to infer transmission without requiring longitudinal sampling or individual strain tracking (*27*–*30*), we asked whether persistence differences could be detected using strain sharing alone. To do so, we returned to our initial 3-month dataset and applied the same criteria we used to classify transmission modes (≥3 cases with a clear majority mode), without imposing constraints based on the temporal pattern of sharing. The persistence of species that were strain-shared within groups (n=39) was not significantly different from the persistence of species shared across groups (n=66) (Cox model, HR=0.987, p=0.930; Figure **S17**). Annotating social transmission therefore requires repeated biological and behavioral sampling over time.

### Fitness implications of the socially transmitted microbiome for hosts

Social transmission of the microbiome has been proposed to influence host health and fitness in two ways. First, social transmission may provide animals with a reservoir or “pan-microbiome” that facilitates microbiome recolonization after disturbances (*5, 31*). Second, socially transmitted species could directly affect host condition (*12*). Using the 18-year metagenomic data set, we evaluated whether transmission mode was related to (i) microbiome similarity to group-mates; (ii) microbiome stability over time; or (iii) host lifespan, the primary predictor of lifetime reproductive success in this population (*32*). Here, we focused on adult females because, as the non-dispersing sex, females had more complete life history data and more extensive metagenomic sampling records (Figure **S18**).

Female baboons that were more socially connected had more similar microbiome compositions to their female group-mates sampled during the same season and year than less socially connected females (linear mixed model of individual social connectedness index [SCI (*33*)], ß=0.010, p=0.014; Figure **4a**). Socially connected females also had more stable microbiomes across consecutive years (linear mixed model of individual SCI, ß=0.017, p=0.009; Figure **4b**). Further, as females declined in social connectedness with age (Figure **S19**), the richness of ST species, but not in NST_within_ or NST_between_, decreased (ST species: linear mixed model, ß=-2.99, p<0.001; NST_within_ species: ß=0.115, p=0.102; NST_between_ species: ß=0.001, p=0.993; Figure **4c**). Individuals who experienced the sharpest decline in their social relationships also experienced the greatest changes in ST species richness (r=0.292, p=0.034; Figure **S19**). Together, these results support the idea that social partners can serve as reservoirs for socially transmitted microbes.

**Figure 4.**
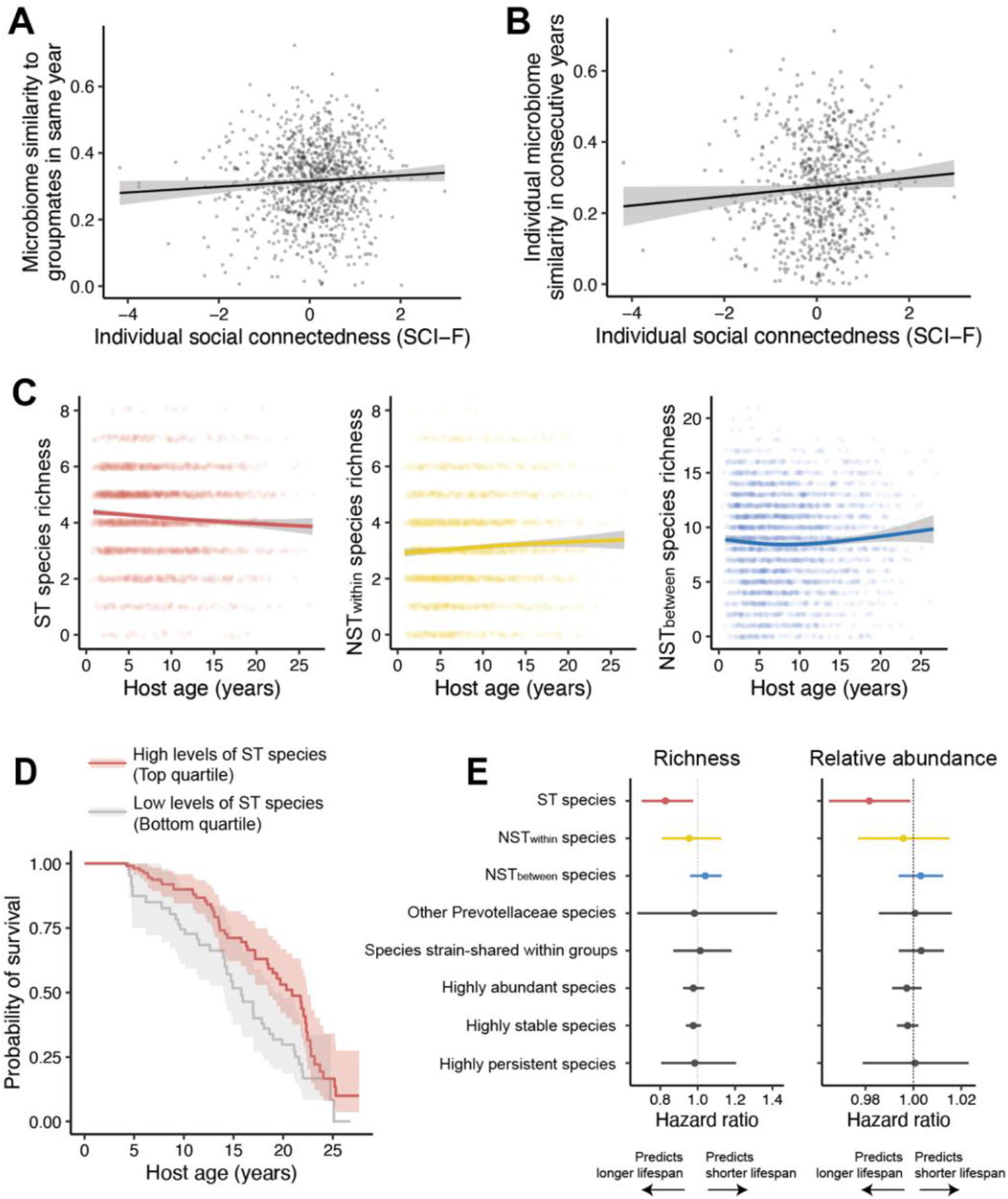
Consequences of social transmission for hosts. **(a)** Individual social connectedness of each female baboon predicted Bray-Curtis similarity across all gut microbial species to other samples from female group members taken in the same season of the same year (ß=0.010, p=0.014). Social connectedness is measured as observer-effort-controlled grooming rates with all other females in her social group, per year of a female’s life. **(b)** Individual social connectedness of each female predicted Bray-Curtis similarity to her own samples taken in the same season of the year before or after (ß=0.017, p=0.009). **(c)** Decrease in ST richness in older baboons (ß=-2.99, p<0.001), but not for NST_within_ or NST_between_ species. **(d)** Female baboons in the upper quartile (5-8 species) of ST species richness live longer on average than females in the lower quartile (0-3 species) (HR=0.827, p=0.024). **(e)** The richness of ST species uniquely predicts female survival. The initial model contained the first four terms (ST richness, NST_within_ richness, NST_between_ richness, Shannon diversity), while subsequent models substituted the ST richness term with the alternative predictor shown. All covariates conformed to the proportional hazards assumption of the model (p>0.05 for all relationships between Schoenfeld residuals and time).

To investigate the functional importance of socially transmitted microbes to survival, we used Cox mixed models with time-varying microbiome measurements. Controlling for other known predictors of adult lifespan (*36*–*39*), both the richness and relative abundance of socially transmitted species positively predicted host lifespan (Cox mixed model, ST richness HR = 0.827 and p=0.024; ST abundance HR = 0.982; p=0.036; Figures **4d, 4e**; Tables **S13, S14**). The average female in the upper quartile of ST species richness is therefore predicted to live 2.5 years longer than a female in the lowest quartile, corresponding to a 12.9% difference in expected lifespan. In contrast, longevity was not predicted by the richness or abundance of NST species, the richness or abundance of non-ST members of the family Prevotellaceae, overall microbiome richness, overall Shannon diversity, or the richness or abundance of species with similar stability, abundance, or persistence as ST species (all p>0.20; Figure **4e**). Indeed, random draws of a same-sized sample of other microbial species in our data set show that few possible sets produce a similar association with lifespan, whether considering all species detected in the population (p=0.049) or the subset implicated in any transmission event (p=0.029; Figure **S20**). Thus, species for which we detected a predominantly social transmission mode are distinctively predictive of longevity.

Because socially connected females generally live longer in this population (*35, 40, 41*), we considered the possibility that the relationship between ST species and survival could contribute to, or be confounded by, social connectedness. However, adding either a composite measure of social relationship quantity (*33*) or of social relationship quality (*42*) did not affect the association between ST species richness and abundance with lifespan. In fact, ST species richness was not correlated with social relationship quantity, and only weakly correlated with social relationship quality (social relationship quantity: ß=0.010, p=0.305; social relationship quality: ß=0.017, p=0.077; Figure **S21**). The lack of a strong relationship between sociality and ST species richness may occur because even the most social individuals are limited to interactions within their social groups. Not all microbial species are found in all groups at all times (Figure **S22**), and ST species were particularly social-group-specific (ANOVA on Bray-Curtis dissimilarities within vs. between groups; ST species: F=83.06, p<0.001; NST_within_ species: F=53.96, p<0.001; NST_between_ species: F=34.87, p<0.001). In other words, while strong social relationships may spread and stabilize ST species within groups, they cannot add species that are not present. Our findings therefore indicate that ST species richness and social relationship quantity and quality, at least based on the measurement approaches used for this population and many other primates, may have independent effects on longevity.

Finally, and similar to our findings for bacterial persistence above, considering strain sharing as the sole criterion for transmission does not recapitulate effects on host lifespan. Neither the richness of species predominantly strain-shared within groups or between groups predicted lifespan (Cox mixed model, HR for species shared within groups=1.040, p=0.297; HR for species shared between groups=0.969, p=0.129; Figure **S23**).

### Social transmission and host-microbe co-evolution

Our results show that some microbial species are primarily transmitted through conspecific social interactions, and that these species lack genes that support survival outside of hosts. Social transmission within species may therefore contribute to host-microbe co-diversification (*6, 7, 43, 44*). If so, ST microbial species should have more narrow host ranges, and strain diversification for ST species should more closely mirror the phylogenies of their hosts.

To test these two predictions, we drew on 1053 publicly available gut metagenomes from 28 phylogenetically diverse primate species, all sampled in the wild (Figure **5a-c**, **S24**). ST microbial species were often detected only in the Amboseli baboons or in other African monkeys (mean number of host species=1.25, mean phylogenetic breadth=3.995 million years). In contrast, NST species had broader host ranges, with some appearing in great apes or even distantly related lemurs (mean number of host species=2.029, mean phylogenetic breadth=20.274 million years; Figure **5d**). ST species were therefore more phylogenetically restricted in the primate radiation (number of host species: p=0.002; phylogenetic breadth: p = 0.016).

**Figure 5.**
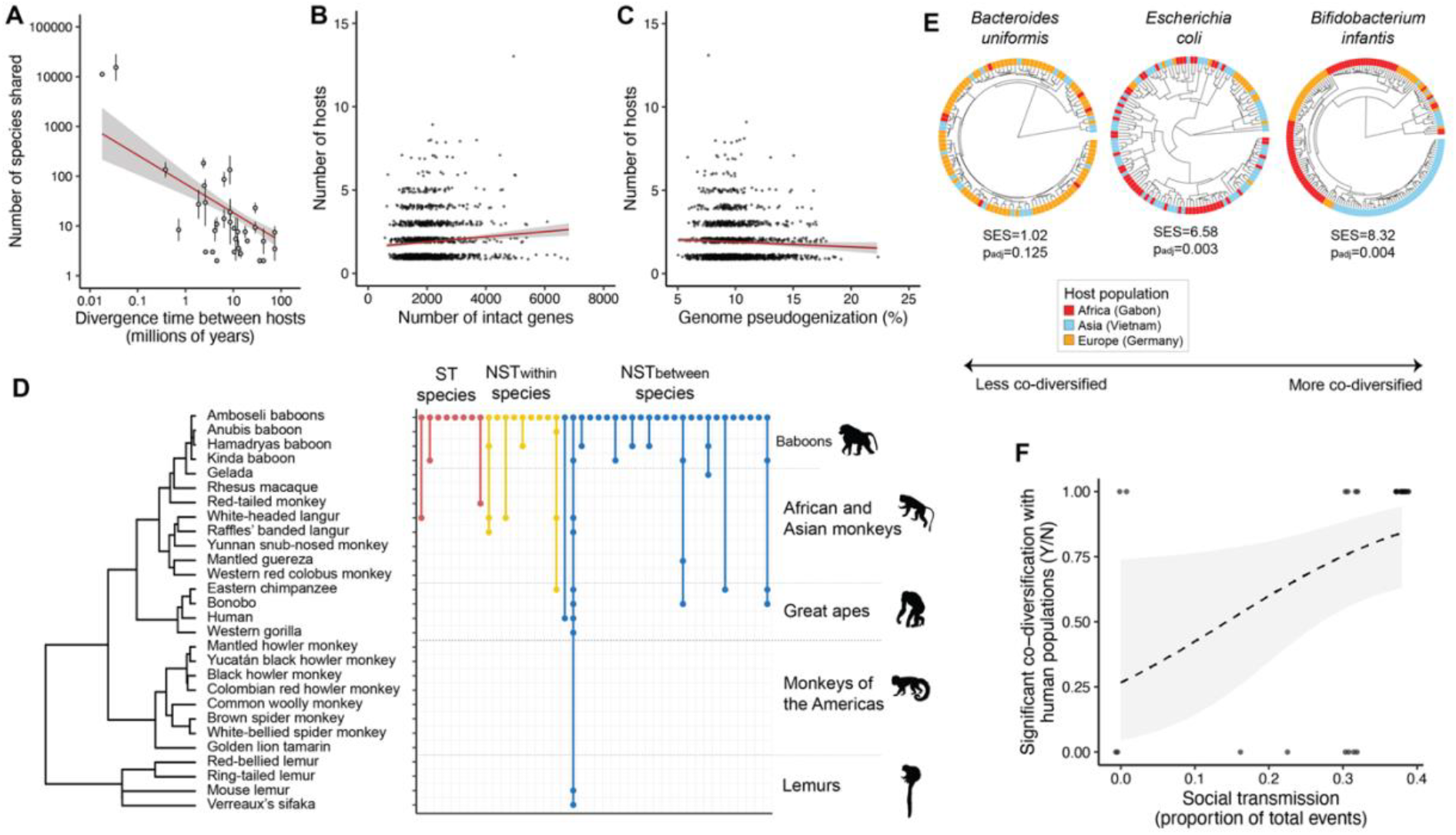
Consequences of social transmission for host-microbe co-evolution. **(a)** In publicly available gut metagenomes from humans and other wild primate populations, the number of shared microbial species between hosts declines with time since divergence (ß=-0.578, p<0.001). Divergence estimates obtained from TimeTree (*61*) and previous work (*45, 46, 62, 63*). **(b)** Across all microbial species in the reference database, the number of hosts in which a species was detected correlates positively with its genome size. **(c)** Across all species in our reference database, the number of hosts in which a species was detected declines with its genome-wide pseudogenization rate. **(d)** Detection of microbial species with annotated transmission modes in the Amboseli baboons and other wild primate populations shows that ST species are more constrained to baboons and their close relatives than NST species. Each column represents a microbial species and lines between species indicate host range. **(e)** Example phylogenies of microbial species detected in human populations from Gabon, Vietnam, and Germany, and associated ParaFit co-diversification statistics (SES=standardized effect size, p_adj_=Benjamini-Hochberg-corrected p-values). **(f)** Human gut species with significant co-diversification population structure come from genera with stronger social transmission patterns, as annotated based on the baboon dataset (ß=7.205, p=0.043).

To test whether social transmission also predicts host-microbe co-diversification in closely related populations, we focused on the three human populations in the comparative dataset, which represented both the densest sampling (100-200 individuals per population in Gabon, Germany, and Vietnam (*6*)) and the most recent radiation (∼30,000-70,000 years (*45, 46*)). To link transmission annotations from baboons to the data from humans, we considered the 58 microbial species found in the human dataset that were close relatives (congeners) of species detected in transmission events in the baboons (Figure **S25**). Both the number of social transmission events and the proportion of total transmission events between social partners in the baboon data set predicted the probability of a significant co-diversification signal in humans (PGLMM on number of social transmission events: ß=0.400, p=0.010; proportion of social transmission events ß=7.205, p=0.043; Figure **5f**). In contrast, non-social transmission events did not significantly predict co-divergence (number of NST_within_ events: ß=0.485, p=0.060; number of NST_between_ events: ß=0.734, p=0.071). Thus, co-diversifying species tend to be close relatives of annotated socially transmitted species, but not non-socially transmitted species.

## DISCUSSION

Tracking microbiome transmission is important for understanding how social behavior affects microbiome acquisition and microbiome-mediated health outcomes, and how transmission in turn shapes microbial evolution (*4, 47*). We confidently identify transmission events and pathways in nature by combining strain-resolved metagenomics with repeated sampling of complete social networks. The resulting picture indicates that a highly non-random set of gut microbial species are primarily transmitted between social partners, predicting both host and microbial fitness. Because strain sharing alone lacked comparable resolution, our results reinforce that some strain sharing among group-mates emerges from processes other than social transmission. Studies of social transmission will therefore be most powerful when they incorporate longitudinal behavioral sampling and strain tracking (*13*).

Our results show that socially transmitted species are ecologically and functionally distinct from microbes that do not rely on host-to-host transmission. These findings mirror observations of vertical transmission: maternally acquired strains are highly persistent in the infant gut, but depleted for traits involved in survival outside of the gut (*26, 48*). Socially transmitted taxa also exhibited stronger signatures of co-diversification across a shallow host radiation, and more restricted host ranges across deeper radiations. Hence, the evolution of socially transmitted microbes appears closely tied to the evolution of their hosts (*49*), either because specialization on one host species reduces fitness in others (*50*), or because socially transmitted species have lost traits required to disperse to new host species.

Our findings raise an important question: given that socially transmitted species predict natural adult lifespan, what are they functionally contributing to their hosts? While the species we identified come from multiple lineages, socially transmitted species were enriched within the genus *Prevotella. Prevotella* shows strong co-diversification in humans and chimpanzees (*6, 51*), and unusually high heritability in baboons (*52*), consistent with a role for social transmission in host-microbe co-evolution. In humans, *Prevotella*-rich gut microbiomes have been linked to glucose homeostasis and weight management (*53, 54*), potentially due to their ability to degrade complex dietary fibers (*51*). Another socially transmitted species belonged to the Christensenellaceae family, which is also involved in complex fiber breakdown (*55, 56*) and has been linked to long lifespan in studies of human centenarians (*57, 58*). Metabolic benefits may thus contribute to the link between social microbiome transmission and lifespan, especially in baboons and humans, which both consume high-starch diets.

Together, our findings support the idea that social microbiome transmission promotes microbial specialization, host fitness, and host-microbe coevolution (*6*). Social transmission may therefore contribute to both microbial specialization and some of the evolutionary benefits of animal social behaviors. However, why some microbes favor social transmission at the apparent expense of more generalist lifestyles remains an open question. Theoretical models find that differences in growth rates, migration rates, and competition between host-associated and free-living environments determine whether selection favors generalism or specialization on the host (*23*) — ideas that annotating microbial transmission modes in natural populations, as we have done here, can now help test. From the host perspective, how animals balance the advantages of acquiring beneficial microbes from social partners with the risks of pathogen exposure is also critical for understanding the evolutionary importance of social transmission. Finally, whether socially transmitted microbes predict morbidity or mortality in other populations, and the extent to which their effects are causal and modifiable, are of direct interest in understanding the ramifications of our findings for human health in rapidly changing social contexts (*59, 60*).

## ACKNOWLEDGMENTS

We thank Jeanne Altmann for her foundational work on the Amboseli baboon population and her foresight in archiving thousands of baboon fecal samples, and Matthew Olm and Hagay Enav for advice on the implementation of strain identification tools. We thank the Core Unit at the Max Planck Institute for Evolutionary Anthropology, the University of Minnesota Genomics Center, and the University of Chicago Genomics Facility for assistance with shotgun metagenomic sequence generation. We also thank the University of Nairobi, the Kenya Institute of Primate Research (KIPRE), the National Museums of Kenya, the members of the Amboseli-Longido pastoralist communities, the Enduimet Wildlife Management Area, Ker & Downey Safaris, Air Kenya, and Safarilink for their cooperation and assistance in the field. Particular thanks go to the Amboseli Baboon Project long-term field team (R.S. Mututua, S. Sayialel, J.K. Warutere, I.L. Siodi, I.L., and L. Musembei), and to T. Wango and V. Oudu for their assistance in Nairobi. The baboon project database, Babase, is managed by J. Gordon and C. Broderick. Database design and programming are provided by K. Pinc.

## Author contributions

R.R.D., E.A.A., and J.T. designed the study. R.R.D. and A.M. collected field samples. R.R.D., C.C.D., S.E.W., P.F., J.G., R.B., E.A.A., and J.T. generated and analyzed sequencing data. S.C.A., E.A.A., and J.T. provided field site administration. R.R.D., E.A.A., and J.T. wrote the manuscript, with revisions from all authors.

## Ethics and inclusion

In Kenya, our research was approved by the Wildlife Research Training Institute (WRTI), Kenya Wildlife Service (KWS), the National Commission for Science, Technology, and Innovation (NACOSTI), and the National Environment Management Authority (NEMA).

## Data, code, and materials availability

The newly generated metagenomic sequencing datasets underlying the analyses in this study have been uploaded to the European Nucleotide Archive (accession numbers PRJEB109125, PRJEB109085, and PRJEB106152). Code used to curate and annotate the reference genome database, map metagenomes to reference genomes, and profile strains and transmission events is available at https://github.com/reenadebray/social-microbiome-transmission.

