## Supplementary Methods and Figures for "Social microbiome transmission predicts microbial specialization and host lifespan in a wild primate"

### Supplementary Information for 'Social microbiome transmission predicts microbial specialization and host lifespan in a wild primate'

Reena Debray, Caroline C. Dickson, Shasta E. Webb, Pamela Ferretti, Alex Meloimet, Jack Gilbert, Susan Alberts, Ran Blekhman, Elizabeth A. Archie, and Jenny Tung

#### TABLE OF CONTENTS


| *Description* | *Page* |
|---|---|
| Materials and Methods | 3 |
| Figure S1: Species-level gut microbiome composition of the Amboseli baboon population. | 12 |
| Figure S2: Integrating SNP-based and synteny-based genome comparison tools. | 13 |
| Figure S3: Strain sharing scenarios that are not considered transmission events. | 14 |
| Figure S4: Detection of transmission as a function of sample availability. | 15 |
| Figure S5: Detection of transmission as a function of sequencing depth. | 16 |
| Figure S6: Mating behaviors do not provide additional insight into drivers of transmission beyond grooming network. | 17 |
| Figure S7: Immigrant males are disproportionately involved in non-social microbial transmission. | 18 |
| Figure S8: Overlap in sleeping site use predicts between-group transmission rates. | 19 |
| Figure S9: Polarization of microbial species towards social or non-social transmission. | 19 |
| Figure S10: Gene content differences between socially and non-socially transmitted species. | 20 |
| Figure S11: Genome size, oxygen tolerance, and temperature tolerance do not differ between socially transmitted and non-socially transmitted species. | 21 |
| Figure S12: No variation in sequencing depth or mapping rates with sample age across 18-year longitudinal dataset. | 22 |
| Figure S13: Abundance and prevalence of socially and non-socially transmitted species in the 18-year dataset. | 23 |
| Figure S14: Seasonal weather in the Amboseli basin. | 24 |
| Figure S15: Microbial persistence within hosts in the 18-year dataset. | 25 |
| Figure S16: Persistence in intensively sampled small cohort. | 26 |

Figure S17: Defining social transmission based on strain sharing within social groups does not identify a social transmission-linked signature of microbial persistence. 26

Figure S18: Female baboons are more densely sampled than males in the 18-year longitudinal metagenomic dataset. 27

Figure S19: Social aging in female baboons. 27

Figure S20: Randomly subsampled pools of species seldom capture survival effect of socially transmitted species. 28

Figure S21: Relationship between social connectedness, distribution of socially transmitted species, and survival. 29

Figure S22: Number of ST species detected per group in each year. 30

Figure S23: Defining social transmission based on strain sharing within social groups does not identify a social transmission-linked signature of host survival. 30

Figure S24: Overview of primate comparative metagenomic dataset. 31

Figure S25: Microbial species used in human co-diversification analysis. 31

References 32

*Supplementary tables:*

Table S1: Sample inventory of three-month shotgun metagenomic dataset.

Table S2: Taxonomic classifications and study accessions of reference genomes used in microbial genome database.

Table S3: Microbial species detected in strain transmission events.

Table S4: Microbial species with annotated dominant transmission modes.

Table S5: Sample inventory of 18-year shotgun metagenomic dataset.

Table S6: Sample inventory of intensively sampled small cohort.

Table S7: Microbial species with median persistence durations exceeding 1 year.

Table S8: Gene annotations relating to motility, chemotaxis, aerotaxis, or flagella.

Table S9: Spore formation genes identified in previous work.

Table S10: Gene annotations classified as components of toxin-antitoxin systems.

Table S11: Gene annotations relating to starch utilization and transport.

Table S12: Publicly available primate gut metagenomic datasets and accession numbers.

Table S13: Full results of Cox proportional hazards model on host lifespan.

#### Materials and Methods:

*Study population and sample collection*

We studied a population of wild baboons inhabiting the semi-arid savannah ecosystem of the Amboseli basin in Kenya. The Amboseli Baboon Research Project collects detailed longitudinal data on the demography, social group membership, home ranges, social relationships, and ecology of baboons in this ecosystem, with continuous, near-daily observations ongoing since 1971(*16*). Fecal samples were collected opportunistically from individually recognized study subjects within a few minutes after the animal defecated and moved away from the sample. Samples were stored in 95-96% ethanol in the Amboseli Baboon Research Project camp for up to two weeks until they could be transported to the University of Nairobi. Samples used for the three-month shotgun metagenomic dataset (Figure **1b**) were transported to the Max Planck Institute for Evolutionary Anthropology in Leipzig, Germany, where they were frozen at -80°C. Samples used for the 18-year shotgun metagenomic dataset and the 652-sample small cohort (Figures **2a**, **S15**) were freeze-dried at the University of Nairobi for dual use in hormone analysis. Briefly, this procedure consisted of 1) evaporating ethanol from samples under a fume hood, 2) cooling tubes for 30 minutes at -20°C, 3) placing tubes in a freeze-dryer (<-50°C, vacuum at 30 millitor for eventual long-term storage at -80°C).

*Strain-resolved metagenomic profiling*

DNA was extracted using the Qiagen PowerSoil Kit. For the 3-month dataset we used the spin-column-based protocol recommended by the manufacturer. For the 18-year dataset and the 652-sample small cohort, we used the 96-well-plate protocol with slight modifications for freeze-dried samples (described in Grieneisen et al.(*52*)). For all data sets, we prepared DNA libraries using the SeqWell purePlex DNA library prep kit and sequenced the libraries on a NovaSeq X using a paired-end, 150 bp read length design. We removed adapters and trimmed reads using Trimmomatic 0.39(*64*), requiring a minimum length of 70 base pairs and a minimum base quality of 20 within a 4-bp sliding window. Reads mapping to the baboon genome (*Panubis1.0*(*65*)) were identified and removed using *bowtie2*(*66*) in paired-end, end-to-end alignment mode. Host-derived reads typically comprised <1% of total metagenomic reads.

As database representation of non-human microbiomes can be low, we mapped reads to a custom database built from the Unified Human Gastrointestinal Genome (UHGG) database and an additional 2,985 metagenome-assembled genomes from non-human primates(*67*). The gut microbial genomes from non-human primates included 48 members of the baboon population analyzed in this study, sampled in July and August 2012. The full set of genomes was dereplicated at the species level (95% average nucleotide identity) using dRep(*68*). Representative genomes were selected based on completeness, contamination, centrality to other genomes in the cluster, and an additional weight to favor non-human primate genomes over those from humans, resulting in a final database of 4,712 microbial genomes (Table **S2**). Taxonomic classifications from the Genome Taxonomy Database were assigned using GTDB-Tk(*69*). Database customization using the additional non-human primate reference genomes improved mapping rates by a median of 11.25% compared to the UHGG database alone (Figure **S1**).

We used two complementary approaches for strain-level population genetic comparisons. *inStrain* assesses strain sharing based on nucleotide identity and is best suited for microbial species that evolve primarily through single-nucleotide polymorphisms, while *SynTracker* assesses strain sharing based on microsynteny and is best suited for species that frequently evolve through small-scale structural rearrangements. To identify instances of strain sharing based on nucleotide identity, we used *bowtie2*(*66*) to map metagenomes to the reference database, then profiled strains using *inStrain*(*20*). Following the developer recommendations, we considered a strain to be present in a pair of samples if at least 25% of its genome was represented with at least 5x coverage in both samples. We considered two samples to share strains if their strains had ≥99.999% average nucleotide identity (popANI). To identify instances of strain sharing based on microsynteny, we generated assemblies using MEGAHIT(*70*), then profiled strains using SynTracker(*21*) with 2.5-kbp regions and 95% BLAST identity. Following the developer recommendations, we considered two samples to share strains if at least 120 regions were represented in both samples with an average pairwise synteny score (APSS) of 0.95.

*Identifying transmission events*

For each microbial species in each pair of individuals, we examined all pairwise comparisons from all timepoints that met the criteria for strain sharing (99.999% popANI and 25% of genome compared using *inStrain*; 0.95 APSS and 120 regions compared using *SynTracker*) to identify when strain sharing was first detected in each member of the dyad. The individual in the strain-sharing dyad who was sampled earlier in time was treated as the potential donor. We then required a minimum of two subsequent samples from the other individual (the candidate recipient): one sample that did not share the strain with the candidate donor, and a second sample where the donor and recipient were first detected to share strains (Figure **1b**). We filtered this set of potential transmission events to consider only those with one possible donor, e.g., strains that were unique to a single individual before their subsequent spread. We note that this method cannot detect cases of “re-transmission”, in which an individual acquires additional cells of a strain that it already has, and may miss highly transient strains that were lost before the recipient was resampled.

As our criteria for strain transmission relied on confidently detecting when strains are shared or not, we considered whether sequencing depth could affect our inferences. Our samples initially ranged from approximately 20-50 million reads/sample, with some as deep as 200 million reads. We included a commercial DNA standard in our sequencing library (ZymoBIOMICS Microbial Community Standard II, Log Distribution), which indicated that we had sufficient depth to detect populations with relative abundance as low as 1 in 10,000,000 cells (Figure **S5a**). There was little relationship between sequencing depth and detection of transmission in general, suggesting that increasingly deep sequencing does not substantially improve our ability to detect transmission, but we noticed a small cluster of samples under ~15 million reads that were never implicated in any transmission events.

We performed targeted resequencing of all samples under 15 million reads to increase their coverage, which resulted in no remaining relationship between coverage and transmission detection (binomial regression, p=0.544, Figure **S5**).

*Social network construction*

Behavioral and demographic data were collected by experienced observers on a near-daily basis, with all groups observed at least twice per week during the period represented by the 3-month dataset (typically 2-4 times). In addition to collecting census data on group membership and reproductive state, observers also routinely collected random-ordered 10-minute focal samples of juveniles and adult females, leading them to move through the group in a pre-determined order while simultaneously recording all grooming interactions that occurred in their line of sight. This approach, known as representative interaction sampling, correlates closely with interaction frequency estimates derived from focal samples alone and provides more even representation of group members(*33*). Because observer effort is more sparsely distributed across group members in large social groups, we regressed the logarithm of observer effort (number of focal samples per female per day) on the logarithm of grooming rate (number of grooming interactions per day) and used the residuals as our measure of grooming frequency. We classified close social partners as within-group dyads with observation-normalized grooming rates in the top 20% of dyads in the population.

*Categorizing transmission events*

Each strain transmission event was annotated as socially transmitted if it occurred between close grooming partners (top 20% of grooming frequencies), transmitted between non-partners if it occurred between members of the same group outside of the top 20% of grooming frequencies, or transmitted across social groups. We then classified microbial species as predominantly transmitted socially (ST); predominantly non-socially transmitted, but between non-partners within the same social group ($NST_{within}$); or predominantly non-socially transmitted between partners that were members of different social groups ($NST_{between}$). Classifications were based on observing at least three transmission events, among which one transmission mode was unambiguously more common than the other two (i.e., no ties).

Importantly, our finding that transmission was enriched among social partners was robust to the interaction frequency we used to define close grooming partners. We report results based on a 20% cut-off in the main text, but a qualitatively similar enrichment is observed whether we define close grooming partners based on the top 10% to 50% of grooming frequencies (top 10% $\log_2$(OR)=0.887; top 30% $\log_2$(OR)=1.253; top 40% $\log_2$(OR)=1.407; top 50% $\log_2$(OR)=1.412; all p<0.001).

Mating-related behavior did not explain additional variation in transmission patterns in this dataset, likely because we detected only two transmission events between mates during this period, and in both cases the pairs were also close grooming partners (Figure **S6**).

*Group movement and home range calculation*

Each group was observed on an alternating schedule that balanced morning and afternoon shifts. Consequently, during the course of a week, each group was followed during most daylight hours. During morning shifts, an observer recorded the location of the sleeping grove in which the baboons had spent the previous night. During afternoon shifts, an observer recorded the location of the sleeping grove in which they ended their day, if the baboons had ascended to a sleeping site before the end of observations. At every half-hour interval, an observer recorded a GPS reading, standing as close as possible to the center of the group without disturbing the animals.

Home ranges were estimated for visualization in Figure **1b** using 95% autocorrelated kernel density estimation (AKDE)(*71*). Home range overlap between two groups was calculated as the Bhattacharyya overlap between their spatial occupancy probability distributions. We assessed significance using a Bayesian dyadic regression model with a Poisson distribution, which incorporates multi-membership random effects. The response matrix was a matrix of transmission events between all group pairs, and the sampling effort matrix was the total number of samples collected from each pair of groups.

$$transmission\ events \sim home\ range\ overlap\ + 1|sampling\ effort$$

To assess overlap in sleeping grove use, we asked how many unique groves had been used by both groups in a group pair at any time in the study period:

$$transmission\ events \sim sleeping\ grove\ overlap\ + 1|sampling\ effort$$

*Genome content and trait predictions*

We used prokka(*72*) to search the 4,712 microbial reference genomes for proteins in the UniProt(*73*), RefSeq(*74*), Pfam(*75*), and TIGRFAMs(*76*) databases. We used pseudofinder(*77*) to identify candidate pseudogenes that were unusually short (<65% of reference gene length), unusually long (>135% of reference gene length), or fragmented (adjacent protein-coding sequences share 50% or more of the same BLAST hits), as well as intergenic sequences that matched amino acid sequences but had no open reading frame, which are thought to represent highly degraded gene remnants(*78*). Microbial species with broad ecological distributions are generally thought to have larger and more intact genomes(*79*, *80*). We tested each of these hypotheses with a linear fixed effects model:

$$genome\ size \sim number\ of\ hosts\ detected$$

$$percent\ of\ genes\ identified\ as\ pseudogenes\ \sim number\ of\ hosts\ detected$$

In the set of 4,712 microbial reference genomes, those found in a broader distribution of primate hosts had more intact genes and a lower rate of pseudogenization, suggesting that these measures are useful indicators of microbial specialists versus generalists (Figure **5b, 5c**).

We tested *a priori* hypotheses about how social transmission could impact the evolution of four microbial traits: motility, sporulation, toxin-antitoxin systems, and starch metabolism. We calculated microbial carriage of motility genes based on the presence of intact (not pseudogenized) genes encoding motility proteins, chemotaxis proteins, aerotaxis proteins, or the assembly, synthesis, or control of flagella as motility genes (Table **S8**). We calculated carriage of sporulation genes based on previous work that identified 100 genes that best distinguished endosporulating from non-spore-forming bacteria (Table **S9**)(*81*). We identified toxin and antitoxin systems by searching for any annotations that contained "toxin", then filtering the full set to keep only those that encoded toxins in toxin-antitoxin systems (i.e., excluding externally secreted toxins, Table **S10**). We calculated microbial carriage of starch utilization genes based on the presence of Sus genes, TonB-dependent transporters, and amylases (Table **S11**). In each case, we assessed significance using a two-sided t-test comparing gene counts between ST and NST species.

We estimated temperature and oxygen requirements using genomeSPOT(*82*), a gene-free model that uses DNA and protein sequence features such as nucleotide content, nucleotide transition rates, amino acid frequencies, hydrophobicity, and residue frequencies. We validated the genome-based oxygen tolerance classifications for all reference genomes with a genus-level assignment using the phenotypic annotations in the Genomes Online Database(*83*) and found 98.5% and 96.3% concordance with genus-level annotations of aerobic and anaerobic metabolism, respectively.

*Persistence in an intensively sampled small cohort*

As consecutive longitudinal samples from the 18-year dataset were collected several months apart, we considered the possibility that drop-outs and recolonization events could be mistake for continuous persistence. We therefore replicated our analysis in a smaller, more densely sampled shotgun metagenomic dataset of 652 fecal samples collected from 9 adult females. In this dataset, each baboon was sampled a median of 71 times, with only 2-4 weeks between samples from the same individual, reducing the potential for undetected drop-outs and recolonization events (Figure **S16**). ST species again persisted for longer durations than NST species (hazard ratio for drop-outs of ST species = 0.341 compared to NST species, p=0.048; Figure **S16**) and were enriched for long-persisters with median durations exceeding 1 year (Fisher's exact test, ST $\log_2$(OR) = 3.812, p=0.046; no long persisters were identified among $NST_{within}$ or $NST_{between}$ species in this dataset).

*Host lifespan*

Because female baboons remain in their natal group throughout their lives, barring full group fissions or fusions, we had complete life histories with confident birth and death (or censorship) dates for all adult females in the survival analysis dataset (n=244, including 113 deaths). We focused on adult survival, conditioning on survival to the age of 4, which is approximately the earliest age when females reach reproductive maturity(*84*). The median lifespan of a female who reached adulthood in this data set was 19.4 years (95% CI 17.2-21.6).

In our analyses of host survival, we controlled for a composite index of early life adversity that is a known, strong predictor of adult female lifespan in this population(*36*). In brief, this index is a count of the following experiences for a young baboon female: 1) maternal loss, 2) a close-in-age younger sibling, 3) early-life drought, 4) large group size, as a measure of realized resource competition, 5) low maternal rank, 6) maternal social isolation. All components were defined based on binarizing the exposures as in previous work(*39*). The other major predictor of adult lifespan known for females in this population is the strength of affiliative social relationships(*42*). We did not include affiliative social relationships (e.g., social bond strength based on the top three bonds with other adult females or adult males, operationalized in the Amboseli baboons as DSI-F and DSI-M(*42*), or social connectedness based on overall grooming to individuals of the same or opposite sex, which has been measured using summaries called SCI-F and SCI-M(*85*); see below) in initial models because they might directly affect the detection of ST species. However, in *post hoc* analyses, including these measures in the survival models did not qualitatively change the relationship between ST richness and lifespan (Figure **S21**).

We estimated microbiome contributions to mortality risk using the following Cox proportional hazard models, focused on richness and abundance, respectively (full model results available in Table **S13** and **S14**):

$$survival \sim ST\ richness + NST_{within}\ richness + NST_{between} richness + Shannon\ diversity + \ early\ life\ adversity\ index + 1|individual$$

$$survival \sim ST\ rel.abundance + NST_{within}\ rel.abundance + NST_{between} rel.abundance\ + Shannon\ diversity + \ early\ life\ adversity\ index + 1|individual$$

To investigate whether the observed effect of socially transmitted species could be driven by other, correlated microbiome phenotypes, we retained the same model structure but substituted the term for ST richness with each of the following covariates, in turn: 1) the richness of all other members of Prevotellaceae in the microbiome, 2) the richness of species identified as more consistently strain-shared within groups than between groups, 3) the richness of all other highly abundant species (mean relative abundance as high or higher than mean relative abundance of socially transmitted species), 4) the richness of all other highly stable species (within-individual coefficient of variation as low or lower than the mean within-individual coefficient of variation of socially transmitted species), 5) the richness of all other highly persistent species (mean persistence as high or higher than the mean persistence of socially transmitted species), 6) the richness of random draws of 8 species from the set of all microbial species detected in the population, 7) the richness of random draws of 8 species from the set of all other microbial species ever implicated in transmission (Figure **S20**).

We substituted the term for ST abundance in the second Cox model with each of the following covariates: 1) the total relative abundance of all other members of Prevotellaceae in the microbiome, 2) the total relative abundance of species strain-shared within groups, 3) the total relative abundance of highly abundant species, 4) the total relative abundance of other highly stable species, 5) the total relative abundance of all other highly persistent species.

*Social connectedness and social bond indices*

We constructed individual- and year-based indices of adult female social connectedness to adult females ($SCI_F$) and adult males ($SCI_M$)(*33*). The social integration value for a female in a given year was calculated as the mean value of her residual from two regressions: (i) the regression between observer effort and daily rates of grooming *given* (to females for $SCI_F$ and to males for $SCI_M$) for all females alive in the population that year, and (ii) the regression between observer effort and daily rates of grooming *received* (from females and males, respectively) for all females alive in the population that year. Observer effort varies due to differences in social group sizes, where observations are distributed more thinly across large groups. We defined observer effort here as the number of focal samples recorded per female per day of observation on the social group, following(*42*).

To evaluate the relationship between $SCI_F$ and microbiome similarity to group members, we calculated for each female-year the average Bray-Curtis dissimilarity to all other samples from female group members taken in the same year and the same season (wet or dry). We implemented the following linear mixed effects model:

$$avg.BC\ dissimilarity \sim SCI_F + 1|individual + 1|year$$

Next, we modified the above model by substituting the response variable with the average Bray-Curtis dissimilarity to female group members based on only the abundances of ST species, based on only $NST_{within}$ species, or based on only $NST_{between}$ species:

$$avg.BC\ dissimilarity\ ST \sim SCI_F + 1|individual + 1|year$$
$$avg.BC\ dissimilarity\ NST_{within} \sim SCI_F + 1|individual + 1|year$$
$$avg.BC\ dissimilarity\ NST_{between} \sim SCI_F + 1|individual + 1|year$$

To evaluate the relationship between $SCI_F$ and microbiome stability, we calculated for each female-year the average Bray-Curtis dissimilarity to all other samples from the same female, taken in the same season, in either the previous year or the following year. If samples were available from the years before and after, we used both. We implemented the following linear mixed effects model:

$$avg.BC\ dissimilarity \sim SCI_F + 1|individual + 1|year$$

We then modified the stability model by substituting the response variable with the average Bray-Curtis dissimilarity to samples from consecutive years based on ST species, $NST_{within}$ species, or $NST_{between}$ species:

$$avg.BC\ dissimilarity\ ST \sim SCI_F + 1|individual + 1|year$$
$$avg.BC\ dissimilarity\ NST_{within} \sim SCI_F + 1|individual + 1|year$$
$$avg.BC\ dissimilarity\ NST_{between} \sim SCI_F + 1|individual + 1|year$$

To measure population-level changes in the microbiome with age, we used the following linear mixed-effects models:

$$ST\ richness \sim age + 1|individual + 1|year$$
$$NST_{within}\ richness \sim age + 1|individual + 1|year$$
$$NST_{between}\ richness \sim age + 1|individual + 1|year$$

To assess the relationship between social connectedness and the rate of microbiome change with age, we focused on females that were well-represented in the microbiome dataset in later life (at least 4 samples after age 10, n=54 females). For each female, we estimated the direction and magnitude of the age coefficient in each of the following within-individual models:

$$ST\ richness \sim age$$
$$NST_{within}\ richness \sim age$$
$$NST_{between}\ richness \sim age$$
$$SCI_F \sim age$$

To measure close social bonds, we constructed individual- and year-based indices of the quality of a female's social relationships with her closest three female ($DSI_F$) and male ($DSI_M$) grooming partners. We required females to be resident in a group for a minimum of 60 days, and considered all potential grooming partners that overlapped in the group with her for a minimum of 1 day. Pairwise dyadic grooming frequencies were corrected for observer effort and standardized using a z-score transformation, separately for female-female and female-male dyads, against all other adult dyads of the same sex composition in the population that year. The dyadic sociality index for a female in a given year was calculated as the mean DSI value between a female and her top three (male for $DSI_M$ or female for $DSI_F$) grooming partners.

*Host range and host-microbe co-diversification*

We identified existing metagenomic datasets from 27 species of wild primates in the European Nucleotide Archive, as well as three human populations from a recent, geographically diverse metagenomic study(*6*, *62*, *86*–*95*) (Figure **S24**). Reads were trimmed with Trimmomatic 0.39 using the parameters described above, then mapped to the set of 4,712 microbial reference genomes with *bowtie2*. For a species to be considered present in a sample, we required at least 50% of a reference genome to be represented with at least 5x coverage and a 92% average nucleotide identity (the expected minimum read recruitment for a genome database dereplicated at 95% ANI(*96*)). We obtained primate divergence time estimates from the TimeTree of Life database (TToL5, accessed 3 February 2025(*61*)). We calculated the phylogenetic breadth of each microbial species using Faith's phylogenetic index, the sum of all branch lengths of host species in which the microbial species was detected(*97*). We assessed significance with a permutation test that randomly shuffled host species labels across the set of primate metagenomic samples and re-calculated the host range and phylogenetic breadth of each ST and NST species with each permutation. The observed differences in mean host range and mean phylogenetic breadth between ST and NST species rarely occurred in permuted data (number of host species permutation test: p=0.002; phylogenetic breadth permutation test: p = 0.016).

We next explored the relationship between transmission and co-diversification among closely related human populations. Though few microbial species were directly shared between humans and baboons, 58 human gut species in samples from Gabon, Germany, and Vietnam(*6*) came from a genus that was implicated in one or more transmission events in our baboon dataset. We therefore assumed that species *related* to species annotated in transmission events in the Amboseli baboon focal population are more likely to exhibit similar transmission modes. For each of these 58 species, we used the *inStrain compare* function to calculate the population average nucleotide identity (popANI) between each pair of host samples containing the species. We constructed a microbial distance matrix based on genome dissimilarity values (1-popANI) and a host distance matrix based on an estimated divergence time of 70,000 years between African and non-African populations, and an estimated divergence time of 36,000 years between European and East Asian populations(*45*, *46*). We calculated the congruence between microbial and host phylogenetic trees using a ParaFit test(*98*), then controlled for multiple testing using Benjamini-Hochberg correction (FDR<0.05). We then asked whether the likelihood of significant co-diversification was explained by the transmission mode ascertained in the baboon dataset (again working under the assumption that transmission mode is more likely to be similar among closely related microbial species than among distantly related species). To do so, we used a phylogenetic linear mixed model (PGLMM) to control for phylogenetic non-independence among microbial species in our dataset:

$$codiversification \sim proportion\ ST + 1|species + 1|phylo.covar.matrix$$

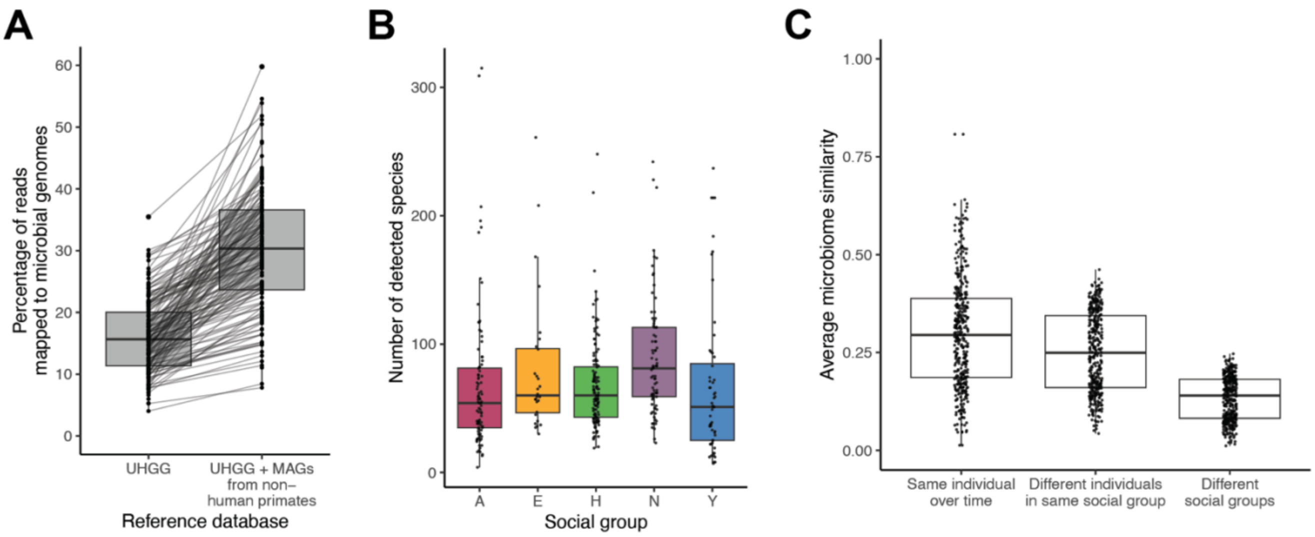


**Figure S1: Species-level gut microbiome composition of the Amboseli baboon population.**

**(a)** Mapping rates of metagenomic reads to standard (Unified Human Gastrointestinal Genome) and custom databases. Adding non-human primate MAGs improves mapping rates by an average of 11.25% (n=410 samples from the concentrated 3-month metagenomic dataset). **(b)** Number of microbial species from our species-representative database of 4,712 microbial genomes detected in each sample, stratified by social group. Group does not explain significant variance in richness (ANOVA, F=1.492, p=0.211). **(c)** Mean microbiome similarity (1-Bray-Curtis dissimilarity) of each sample to other samples from the same individual collected at different times, to samples from different individuals in the same social group, and to samples from individuals in different social groups at any time during the study (Tukey HSD, $p_{adj}<0.001$ for all comparisons)

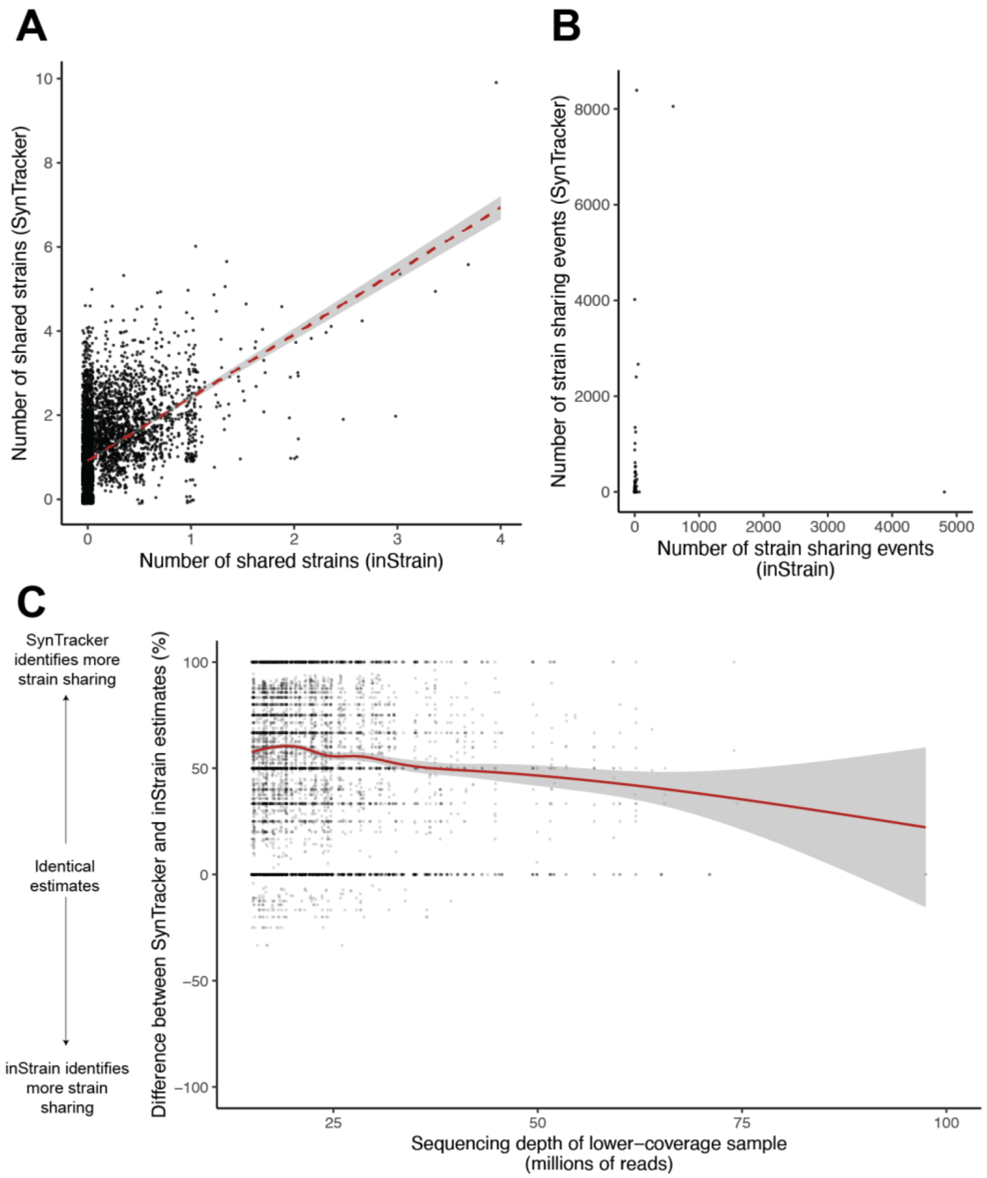


**Figure S2: Integrating SNP-based and synteny-based genome comparison tools.**
**(a)** Dyad-wise estimates of strain sharing by *inStrain* and *SynTracker* are positively correlated (each point represents a pair of samples; r=0.471, p<0.001). Sample dyads that shared more strains based on *inStrain* also shared more strains based on *SynTracker*. **(b)** Species-wise estimates of strain sharing between pipelines are not correlated (each point represents a species; r=0.076, p=0.128), meaning that the two strain-sharing detection approaches are best suited for different microbial species, as expected based on their original design. **(c)** *SynTracker* generally identified more instances of strain sharing per sample pair than *inStrain*, with differences particularly pronounced for lower-coverage samples. The y-axis represents differences in percentage values. For example, a dyad in which 55% of shared species include a shared strain according to *SynTracker* but 40% of shared species include a shared strain according to *inStrain* would have a difference of 15%.

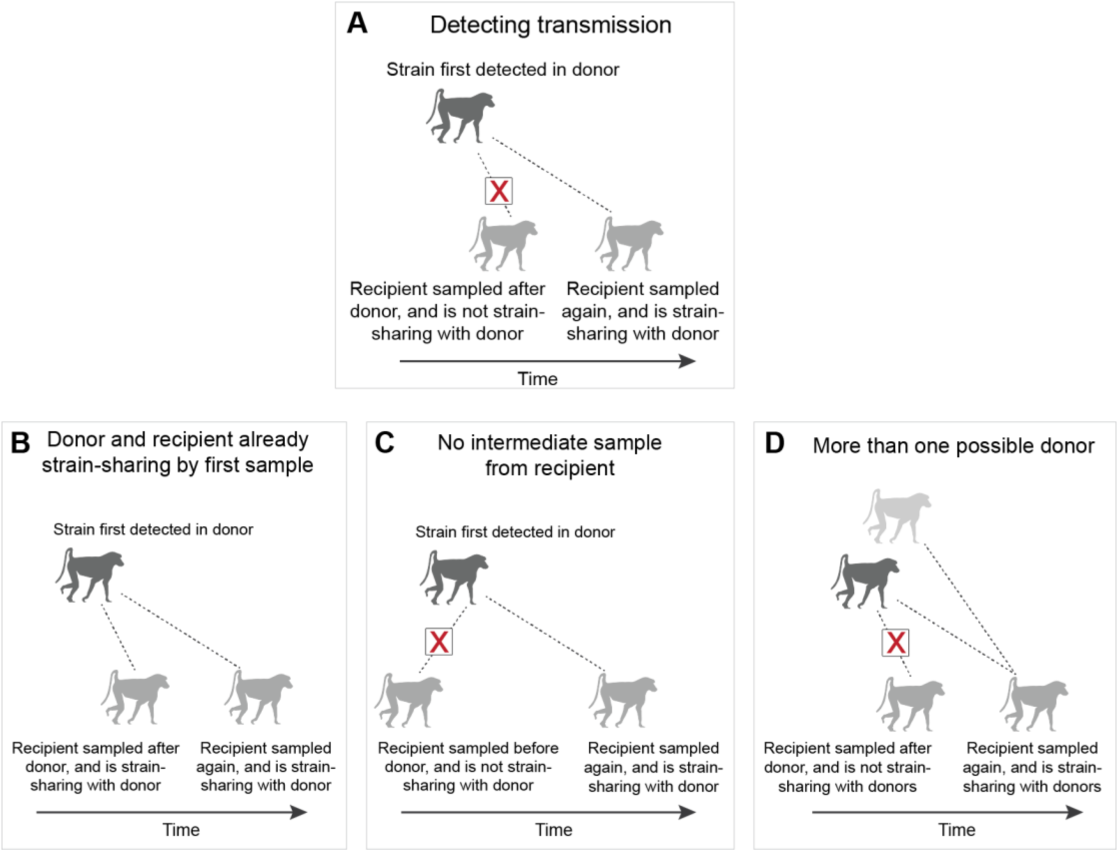


**Figure S3: Strain sharing scenarios that are not considered transmission events.**

**(a)** The scenario *required* to detect transmission (reproduced here from Figure **1c inset** as a reminder to the reader). **(b-c) Other scenarios that are not considered transmission events. (b)** Donor and recipient were strain-sharing in both of their first samples, making it impossible to determine timing or order of acquisition. **(c)** Recipient was not sampled between the time of first detection in the donor and the time of first detection in the recipient, again making it impossible to determine order of acquisition. **(d)** Strain sharing occurs with multiple possible donors (e.g., strain is already widespread in the population), making the timing and order of acquisition unclear, including whether the strain originated from a completely separate individual.

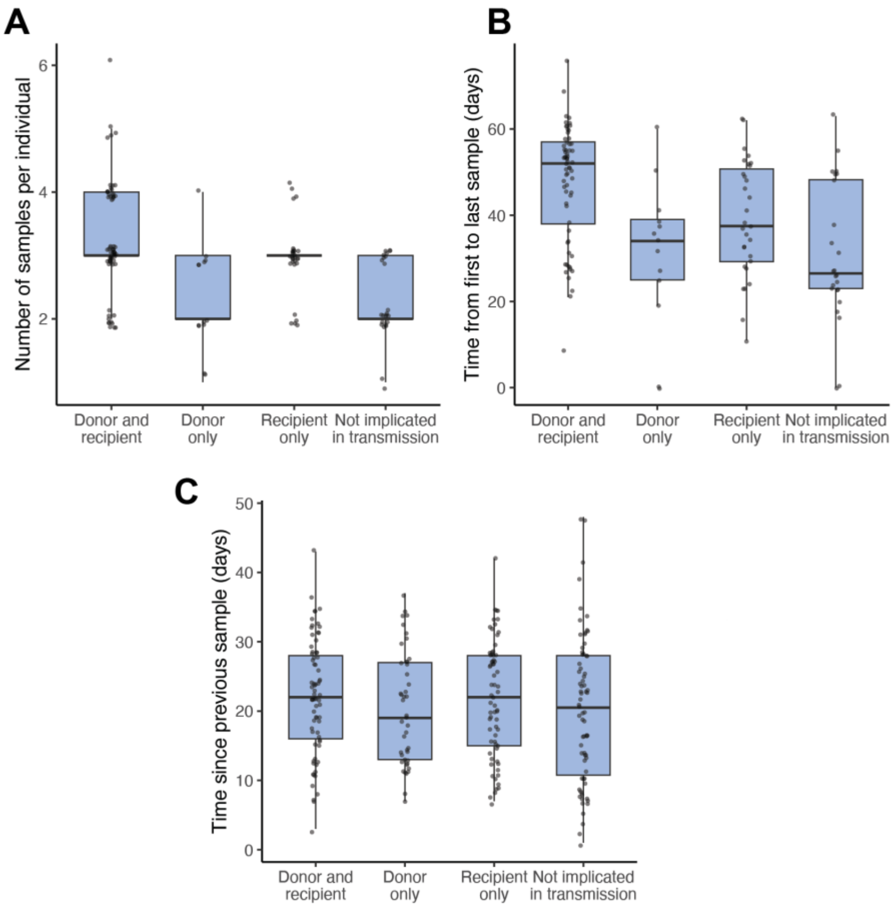


**Figure S4: Detection of transmission as a function of sampling effort per individual.** **(a)** Extensively sampled individuals are more likely to be detected as donors and recipients of transmission (significant comparisons from Tukey HSD test: “Not implicated in transmission”-“Donor and recipient” ($p_{adj}$=0.027) and “Donor only”-“Donor and recipient” ($p_{adj}$=0.023). Dots are jittered to avoid overplotting (all are integer values). **(b)** Individuals sampled over a longer time period are more likely to be detected as donors and recipients of transmission (significant comparisons from Tukey HSD test: “Not implicated in transmission”-“Donor and recipient” ($p_{adj}$<0.001), “Donor only”-“Donor and recipient” ($p_{adj}$=0.001), “Recipient only”-“Donor and recipient” ($p_{adj}$=0.05). **(c)** Samples taken closer together in time are not significantly more likely to be detected as donors or recipients.

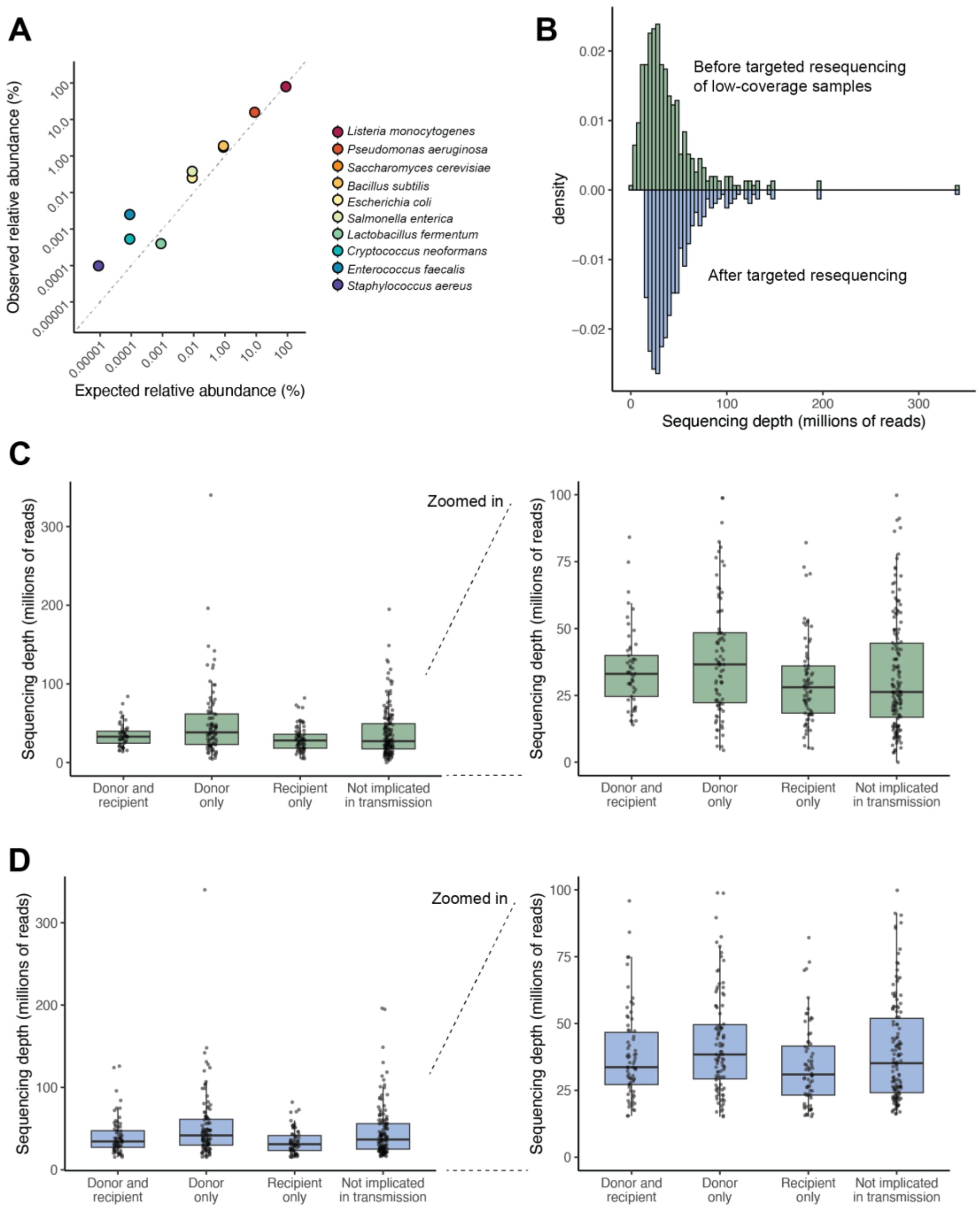


**Figure S5: Detection of transmission as a function of sequencing depth.**
**(a)** Expected versus observed relative abundance of species in a known microbial community (Zymo Microbial Community Standard II, Log Distribution). **(b)** Distribution of read depths before (green) and after (blue) a targeted resequencing effort of all samples below 15 million reads/sample. **(c)** Coverage of samples implicated/not implicated in transmission before resequencing. **(d)** Coverage of samples implicated/not implicated in transmission after resequencing. In (c) and (d), the right-hand plots show the same data as the left-hand plots but are zoomed in to focus on the range of sequencing depths that include nearly all samples (except for a few very deeply sequenced outliers).

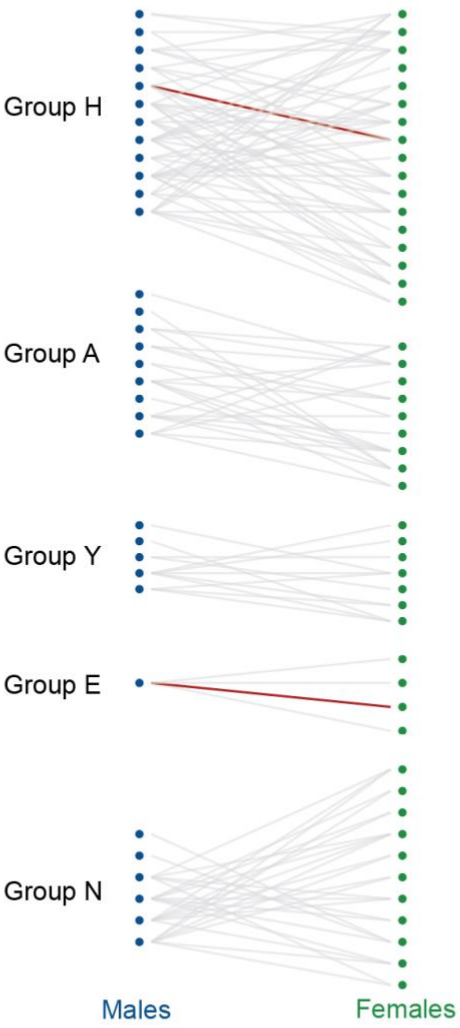


**Figure S6: Mating behaviors do not provide additional insight into drivers of transmission beyond grooming network.**

All consortships (i.e., close associations between adult males and females in oestrus) recorded between July-December 2023 (grey), with transmission between consorting pairs highlighted in red. Both consorting pairs implicated in transmission were also close grooming partners; thus, including information on consortships did not add information about social transmission to our dataset.

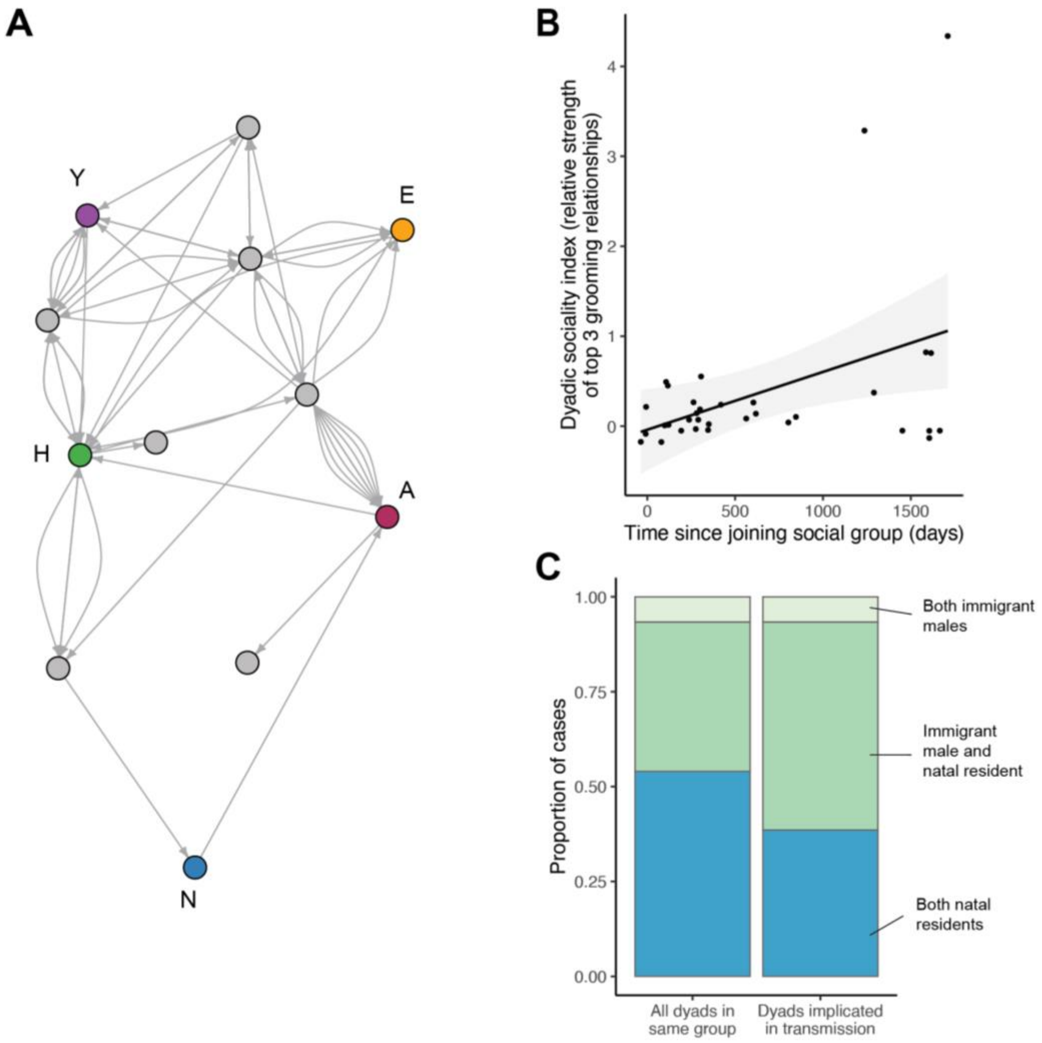


**Figure S7: Immigrant males are disproportionately involved in non-social microbial transmission.**

**(a)** Known dispersal events for males in the 3-month transmission data set prior to sampling, including both dispersals from their natal group at reproductive maturity and subsequent (secondary) dispersals later in life. Colors and labels indicate currently monitored groups (as in Figure **1**); grey nodes indicate other social groups in the Amboseli region. Arrows point from group of residency before dispersal to group of residency after dispersal. The immigrant males in our study include 26 males who dispersed between known groups and 7 males of unknown origin (not plotted in panel **(a)**) who joined their study groups at the time of sampling as adults. **(b)** Social bond strength of immigrant males is initially low after joining a new group, but generally increases with residency time. The y-axis shows a composite measure of the relationship of males to their top 3 female grooming partners (i.e., $DSI_F$). **(c)** Immigrant males are disproportionately involved in detectable transmission events compared to baboons living in their natal group (females or natal males who have not yet dispersed) (Fisher's exact test, $p<0.001$).

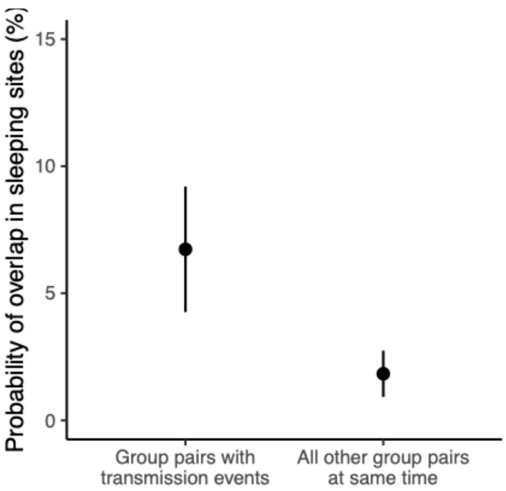


**Figure S8: Overlap in sleeping site use predicts between-group transmission rates.**
For each transmission event, group pairs were considered to overlap in sleeping sites if they used any of the same sleeping groves in the two weeks prior to when strain-sharing was first detected in the recipient. Group pairs associated with each transmission event (n=17, left) were compared to all other group pairs that did *not* experience any transmission events during the same period (dyadic regression, p<0.001).

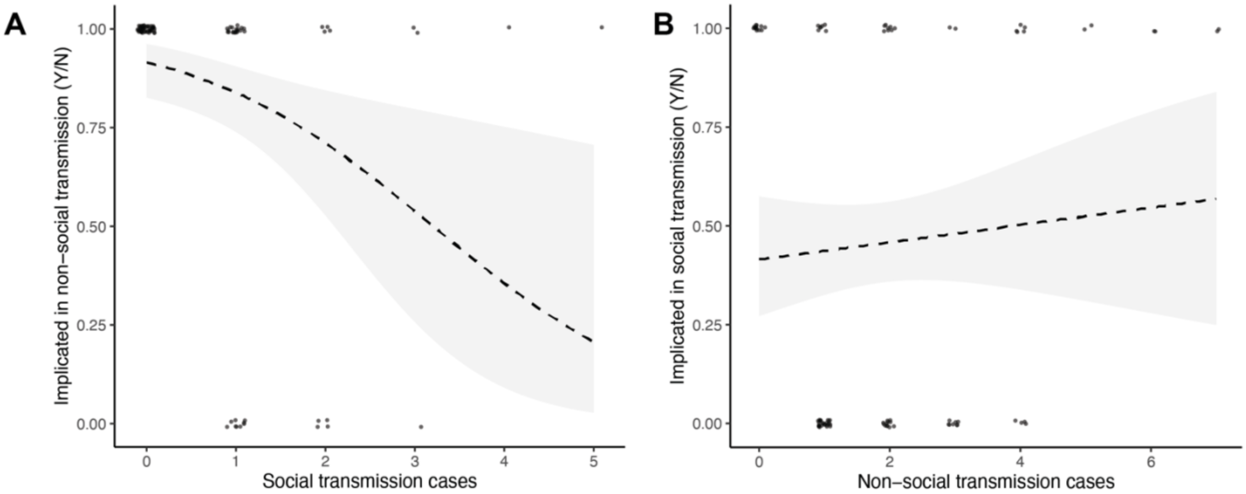


**Figure S9: Polarization of microbial species towards social or non-social transmission.**
**(a)** Species where more social transmission events were identified in our sample were less likely to be implicated in non-social transmission (modeled as a binary outcome: 1 = detected in non-social transmission at least once;  0 = never detected in non-social transmission, ß= -0.747, p=0.007). **(b)** Species with more non-social transmission events were not more or less likely to be implicated in social transmission (modeled as binary outcome: 1 = detected in social transmission at least once; 0 = never detected in social transmission, ß= 0.088, p=0.504).

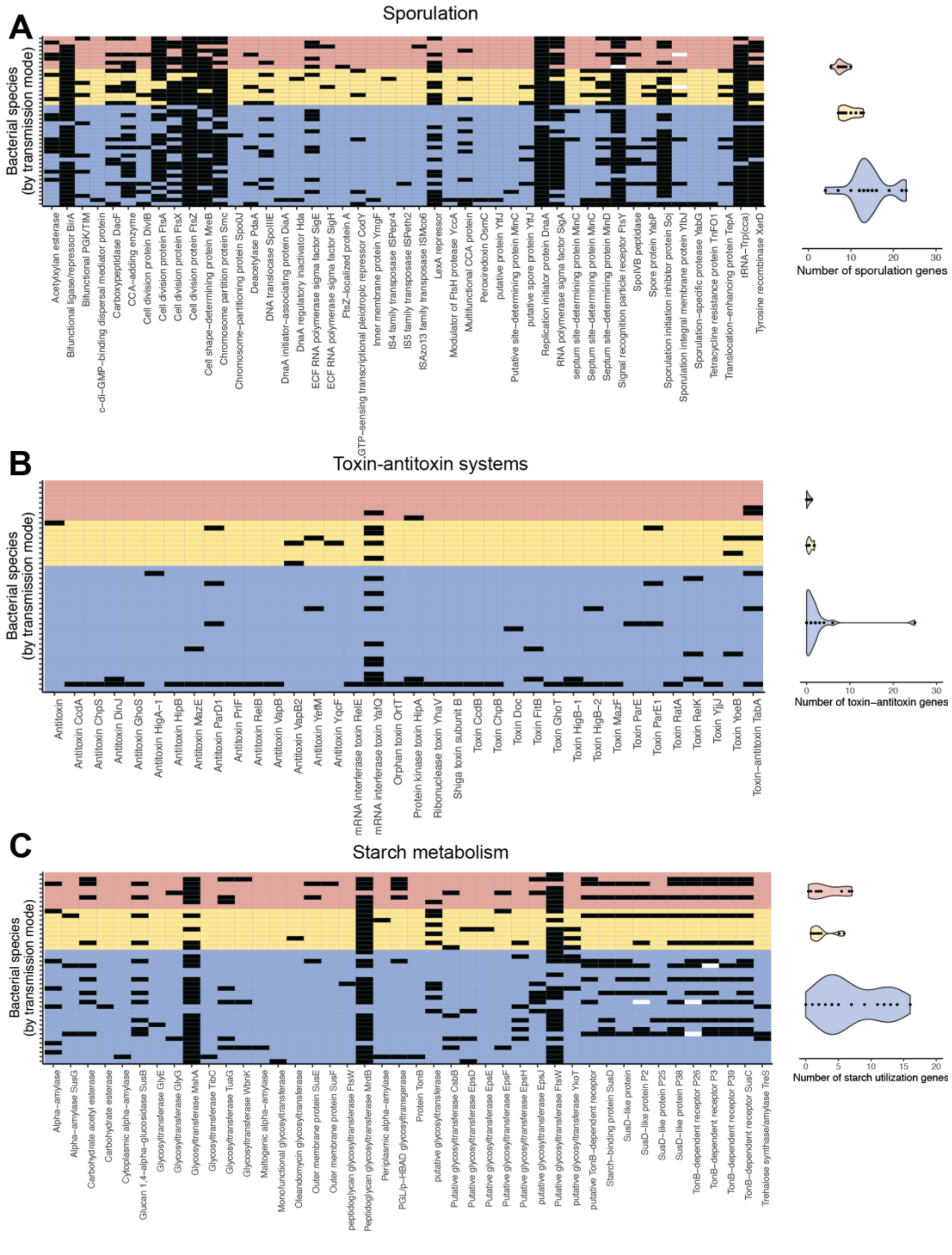


**Figure S10: Sporulation, toxin-antitoxin, and starch utilization genes in the genomes of socially and non-socially transmitted species.**

**(a)** Socially transmitted species carried marginally fewer sporulation genes than NST species (t-test, t=1.804, p=0.086). **(b)** Socially transmitted species carried marginally fewer genes involved in toxin-antitoxin systems than NST species (t=1.92, p=0.063). **(c)** No difference in the distribution of starch utilization genes, TonB-dependent transporters, and amylases (t=-0.550, p=0.596). Black tiles indicate intact genes and white tiles indicate pseudogenes.

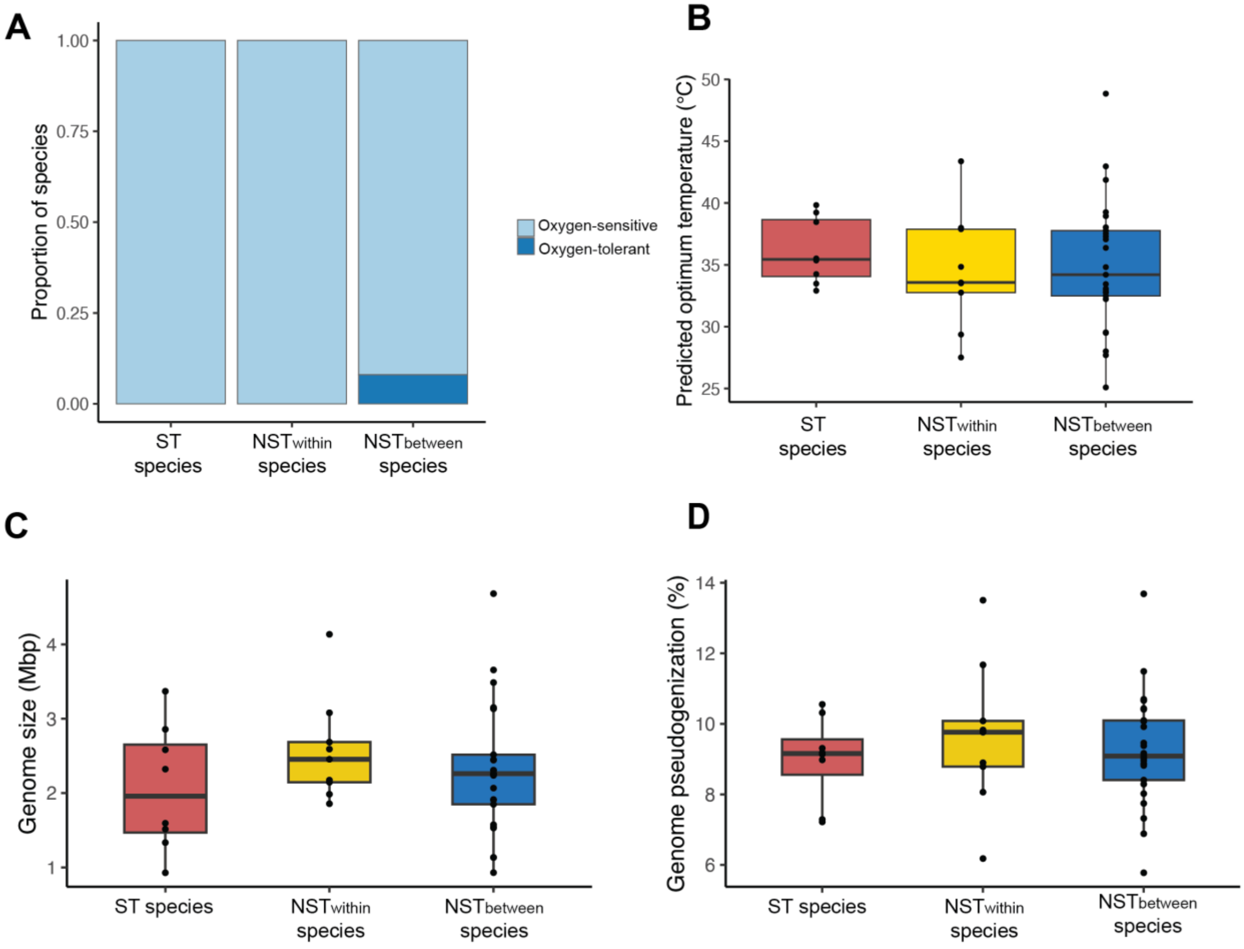


**Figure S11. Oxygen tolerance, temperature tolerance, and genome structure do not differ between socially transmitted and non-socially transmitted species.**
**(a)** No difference in predicted oxygen tolerance by transmission mode based on GenomeSpot annotations (Fisher's exact test, $log_2(OR)=1$, $p=1$). **(b)** No difference in predicted temperature tolerance by transmission mode detected in this population based on GenomeSpot annotations (ANOVA, F=0.0073, p=0.933). **(c)** Total genome size did not vary with transmission mode (ANOVA, F=0.7838, p=0.464). **(d)** The percentage of genes in the genome that were annotated as pseudogenes based on length relative to reference genes, fragmentation, or missing start codons did not vary with transmission mode (ANOVA, F=0.334, p=0.718).

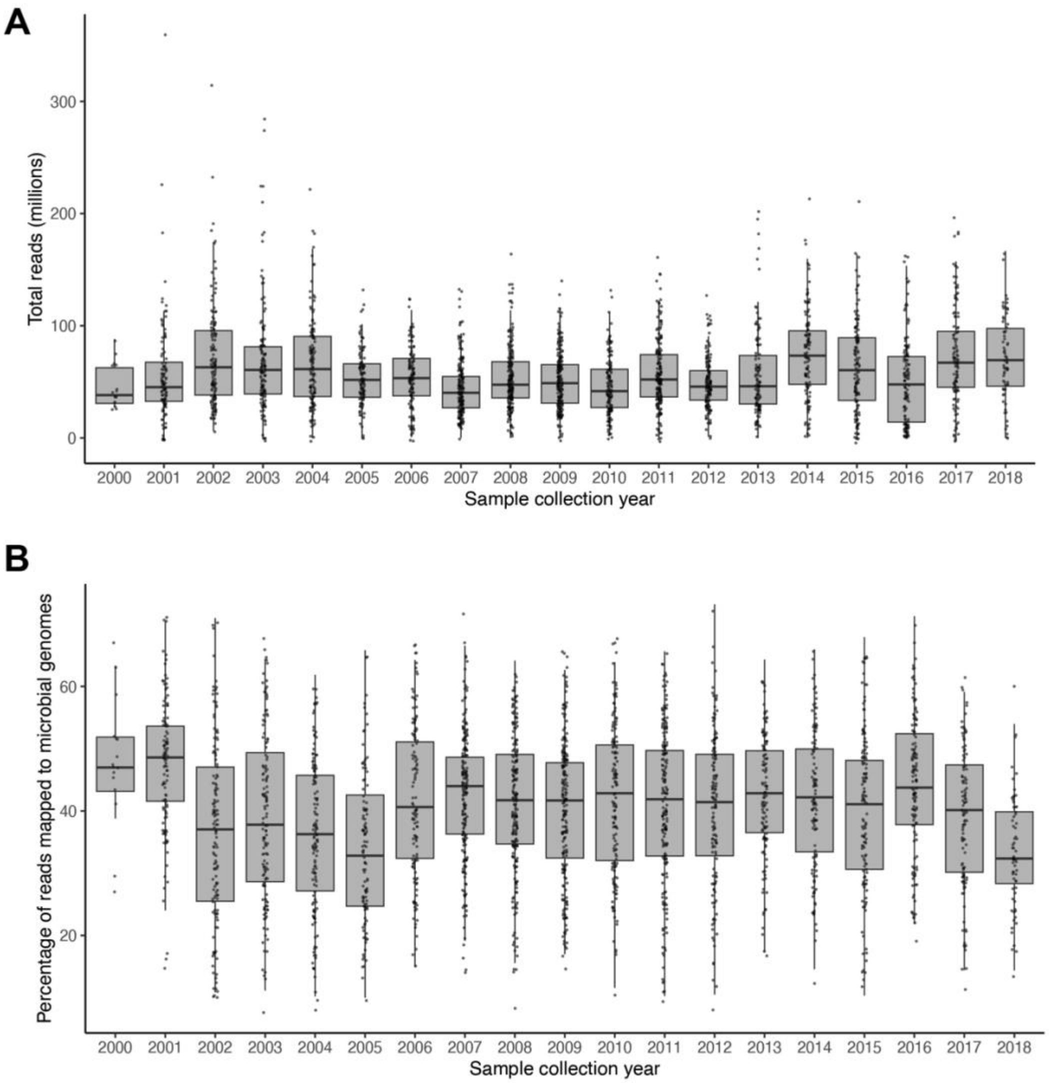


**Figure S12: No systematic variation in sequencing depth or mapping rates with sample age across 18-year longitudinal dataset.**

**(a)** Distribution of sequencing depths by sampling year. **(b)** Read mapping rates to our custom microbial genome database (n=4,712 species-representative bacterial and archaeal genomes) by sampling year.

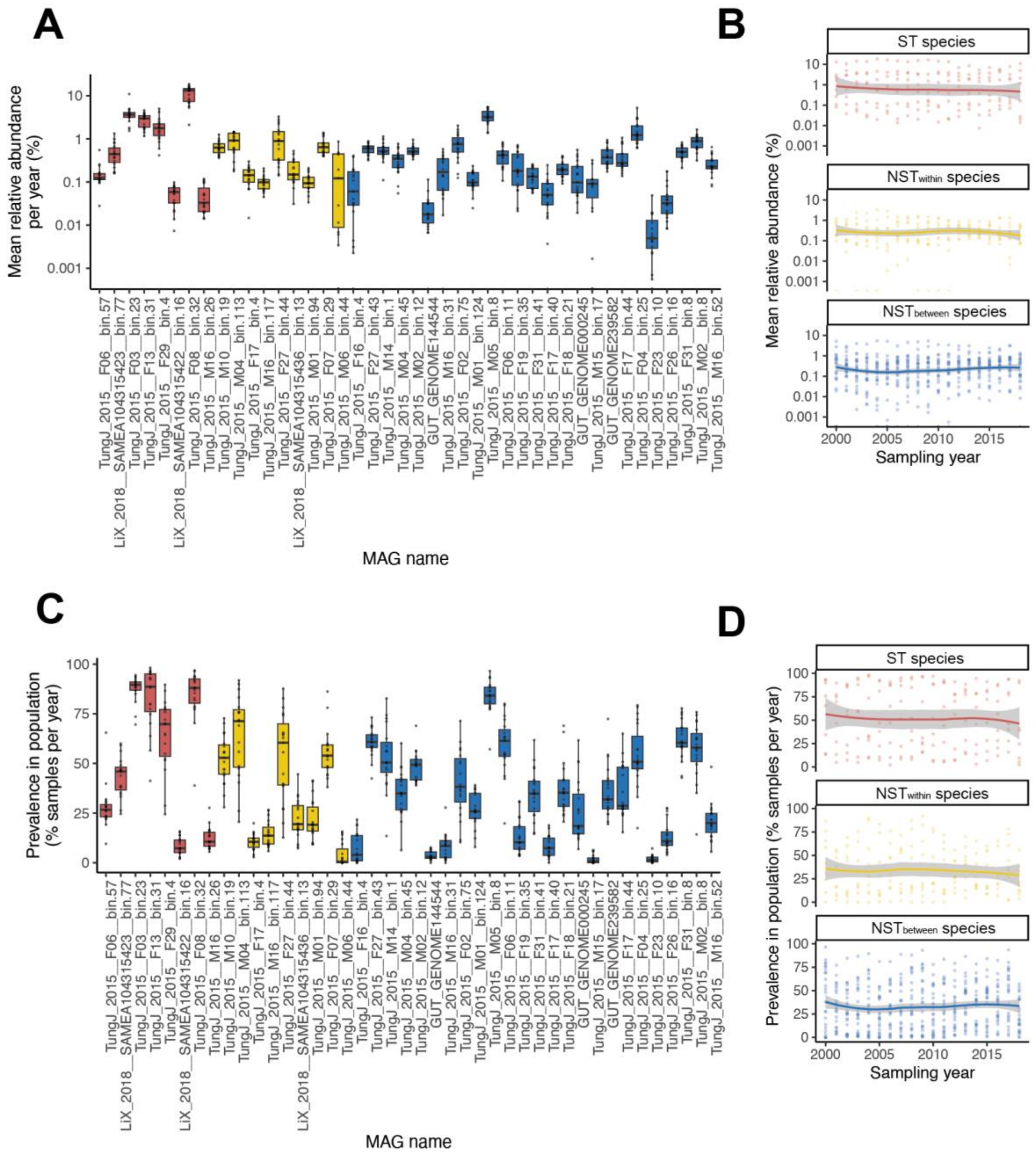


**Figure S13: Abundance and prevalence of socially and non-socially transmitted species in the 18 year dataset.**

**(a)** Within-host relative abundances of ST and NST species (red = ST species, gold = $NST_{within}$ species, blue = $NST_{between}$ species). **(b)** Slight declines in relative abundances of ST and $NST_{within}$ species over time (linear mixed-effects model, ST species: ß= -0.0003, p=0.016; $NST_{within}$ species: ß= -0.00004, p=0.038; $NST_{between}$ species: -0.000007, p=0.612). **(c)** Population prevalence of ST and NST species. **(d)** No significant change in population prevalence of ST and NST species over time (ST species: ß= -0.0023, p=0.174; $NST_{within}$ species: ß= -0.0019, p=0.291; $NST_{beween}$ species: ß=0.0015, p=0.094). All NST and ST species identified in the 3-month dataset used to annotate transmission mode were also detected in the 18-year longitudinal dataset.

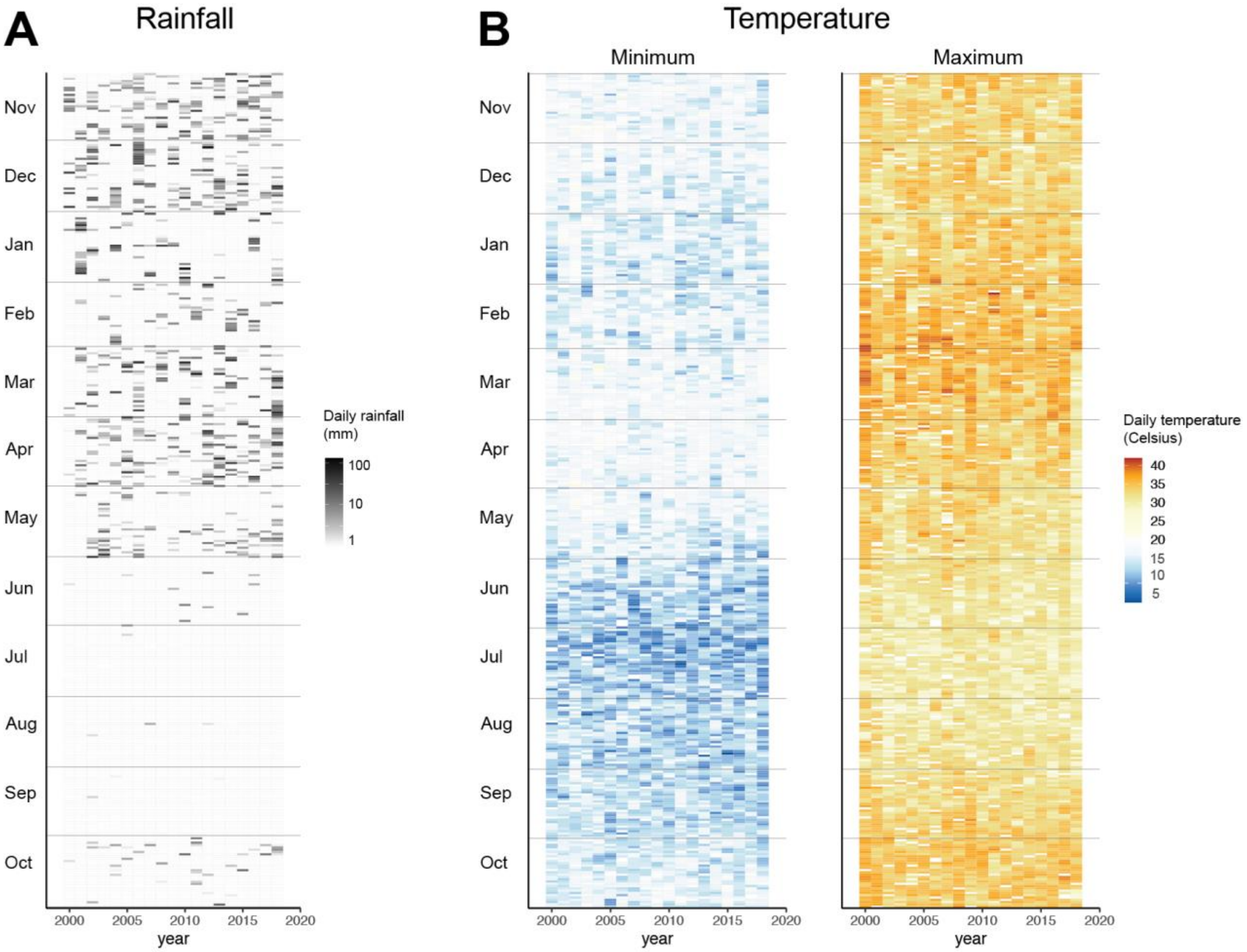


**Figure S14: Seasonal weather in the Amboseli ecosystem.**
**(a)** Daily rain gauge measurements in Amboseli from 2000-2018 across the Amboseli hydrological year (Nov 1- October 31). **(b)** Average daily minimum and maximum temperatures in Amboseli from 2000-2018.

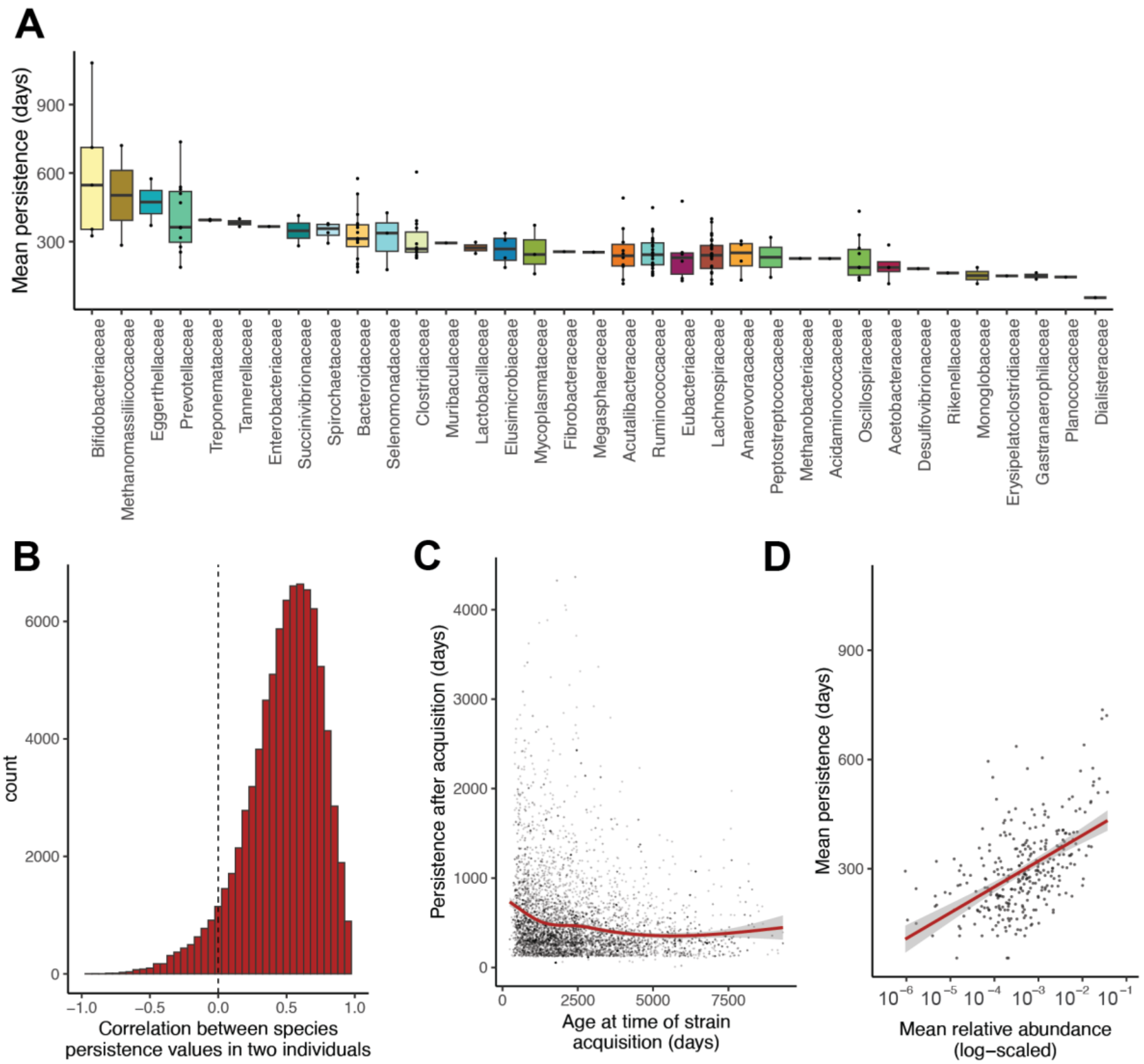


**Figure S15: Microbial persistence within hosts in the 18-year dataset.**
**(a)** Maximum duration of within-individual persistence, averaged across individuals for each microbial species and grouped by microbial family membership. **(b)** Persistence of microbial species was positively correlated across individuals. Each correlation coefficient is based on the maximum persistence of all microbial species shared between two individuals. The majority (94.1%) of pairwise correlations were positive. **(c)** Persistence was elevated for strains acquired early in life (quadratic coefficient=3476.82, $p<0.001$, linear coefficient=-6423.04, $p<0.001$). **(d)** Mean persistence per species was positively associated with mean relative abundance ($r=0.594$, $p<0.001$).

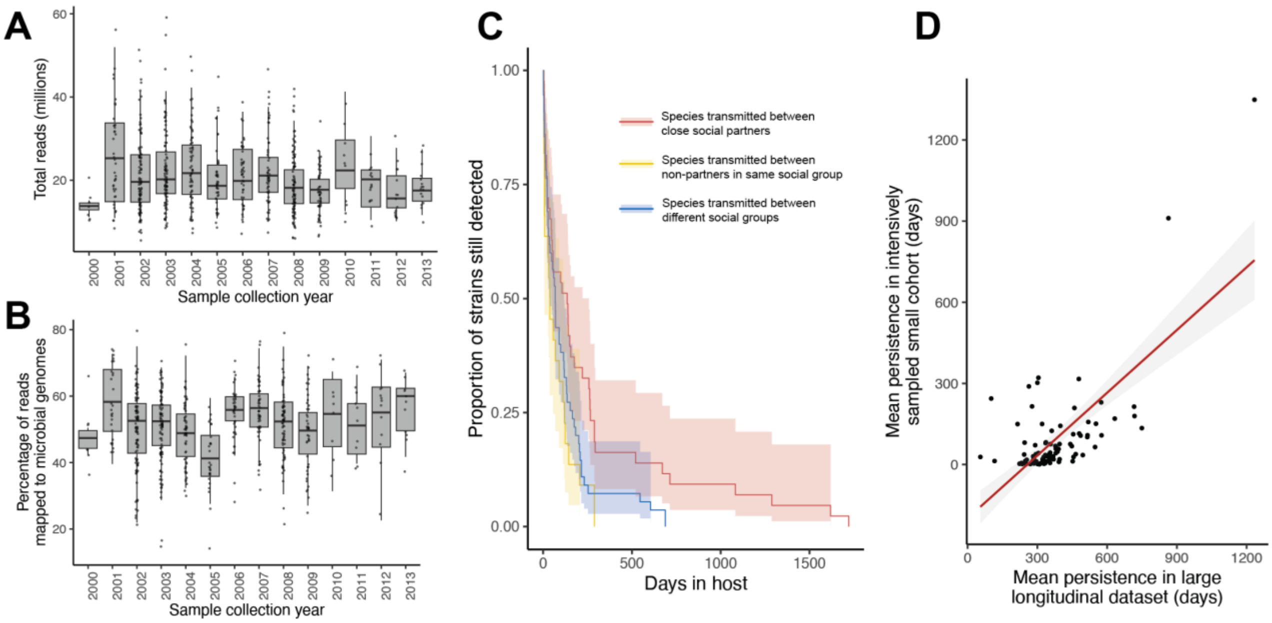


**Figure S16: Persistence in an intensively sampled small cohort.**
**(a)** Distribution of sequencing depths by sampling year in a data set with 652 fecal samples collected from nine adult female baboons. **(b)** Reads mapped to our custom reference database of 4,712 microbial genomes by year of sample collection. Of the 42 ST and NST species identified in the 3-month dataset, 27 were also detected in this small cohort (6 of 8 of ST species, 6 of 9 of $NST_{within}$ species, and 15 of 25 $NST_{between}$ species). **(c)** As in the larger, but more sparsely sampled gut metagenomic data set, socially transmitted species persist for longer durations than non-socially transmitted species (hazard ratio=0.341, p=0.048). **(d)** Average persistence durations per microbial species are correlated across between the small cohort (652 sample/9 individual) dataset and the 18 year dataset (r=0.698, p<0.001).

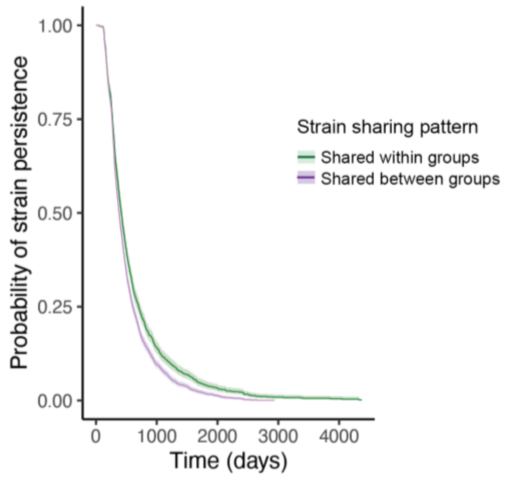


**Figure S17: Defining social transmission based on strain sharing within social groups does not identify a social transmission-linked signature of microbial persistence.**
Persistence of species for which strain-sharing was detected predominantly within groups (n=39 species) did not differ from persistence of those predominantly shared between groups (n=66 species) in the 18-year longitudinal dataset (HR=0.987, p=0.930).

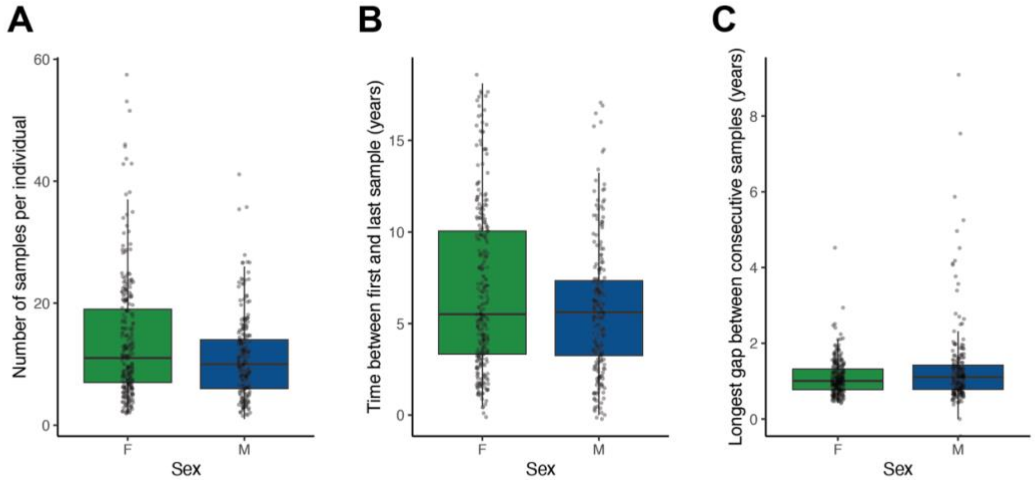


**Figure S18: Female baboons are more densely sampled than males in the 18-year longitudinal metagenomic dataset.**

**(a)** Number of samples per individual differed by sex (ANOVA, F=8.657, p=0.0034). **(b)** Time between first and last sample of each individual differed by sex (ANOVA, F=5.843, p=0.0151). **(c)** Longest gap between consecutive samples from the same individual differed by sex (ANOVA, F=10.704, p=0.0011).

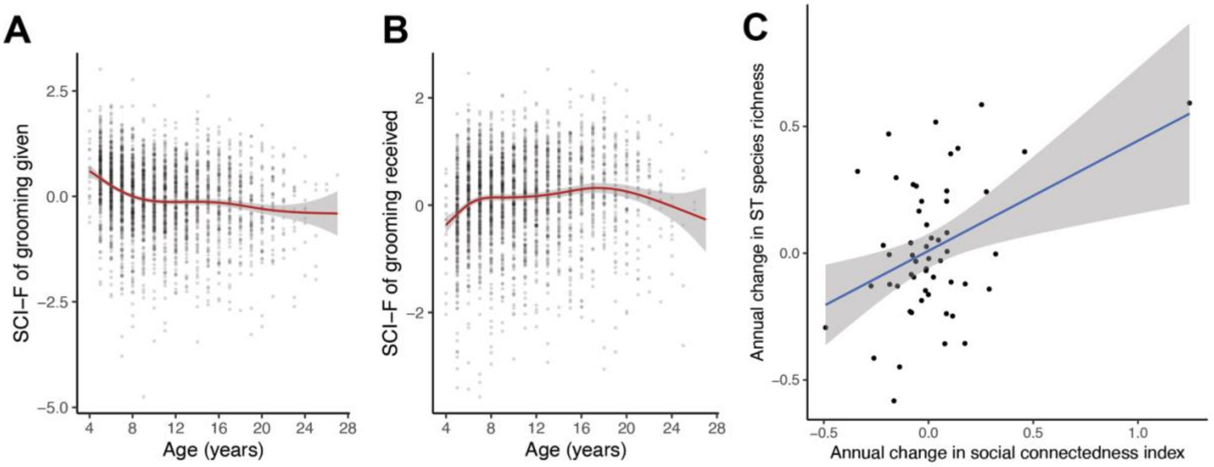


**Figure S19: Social aging in female baboons.**

**(a)** Total grooming given by each female baboon in each year of her life, normalized for observer effort, plotted by her age at the beginning of the interval. Grooming given exhibited a concave up, decreasing relationship with age (linear mixed-effects model controlling for individual, quadratic age coefficient: ß=6.709, p<0.001, linear age coefficient: ß=-6.340, p<0.001) **(b)** Total grooming received by each female baboon in each year of her life. Grooming given exhibited a concave down, decreasing relationship with age (quadratic age coefficient: ß= -13.151, p<0.001, linear age coefficient: ß= 5.021, p<0.001). **(c)** Females with the sharpest declines in social relationships experience the sharpest declines in their social microbiome richness (r=0.292, p=0.034; without upper right outlier: r=0.265, p=0.056). Each point indicates the regression coefficient for social connectedness on age (x-axis) and ST microbial richness on age (y-axis) for all samples collected from an individual female baboon after age 10. Females were included in this plot if at least 4 samples after age 10 were available.

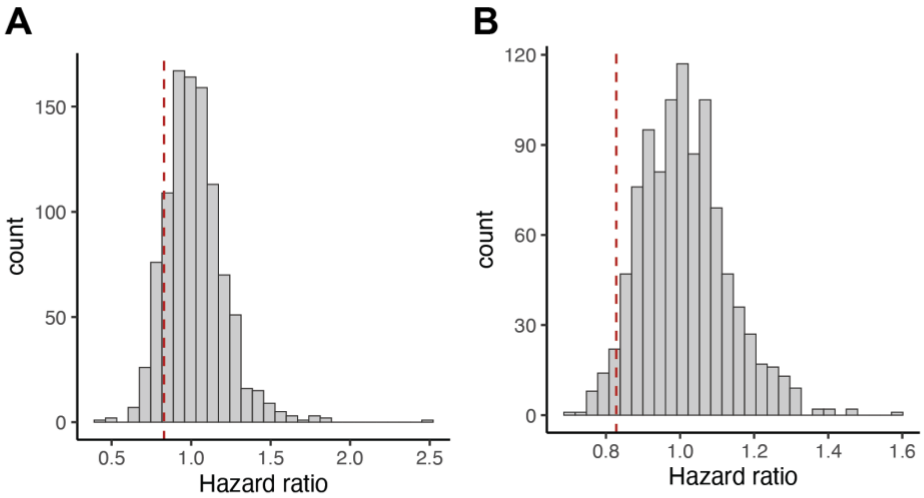


**Figure S20: Randomly subsampled pools of species seldom recapitulate the relationship between the richness of socially transmitted species and female lifespan.**

**(a)** Hazard ratios from models based on 1000 randomly subsampled species pools drawn from all microbial species detected in population (not filtered for detection in transmission events) were rarely as extreme as the observed effect of ST species richness (Hazard ratios: 123 iterations out of 1000, p-values (not shown): 17 iterations out of 1000). **(b)** Hazard ratios from models based on 1000 randomly subsampled species pools drawn from all species with annotated transmission modes, excluding ST species, were rarely as extreme as the observed effect of ST species richness (Hazard ratios: 38 iterations out of 1000, p-values (not shown): 29 iterations out of 1000).

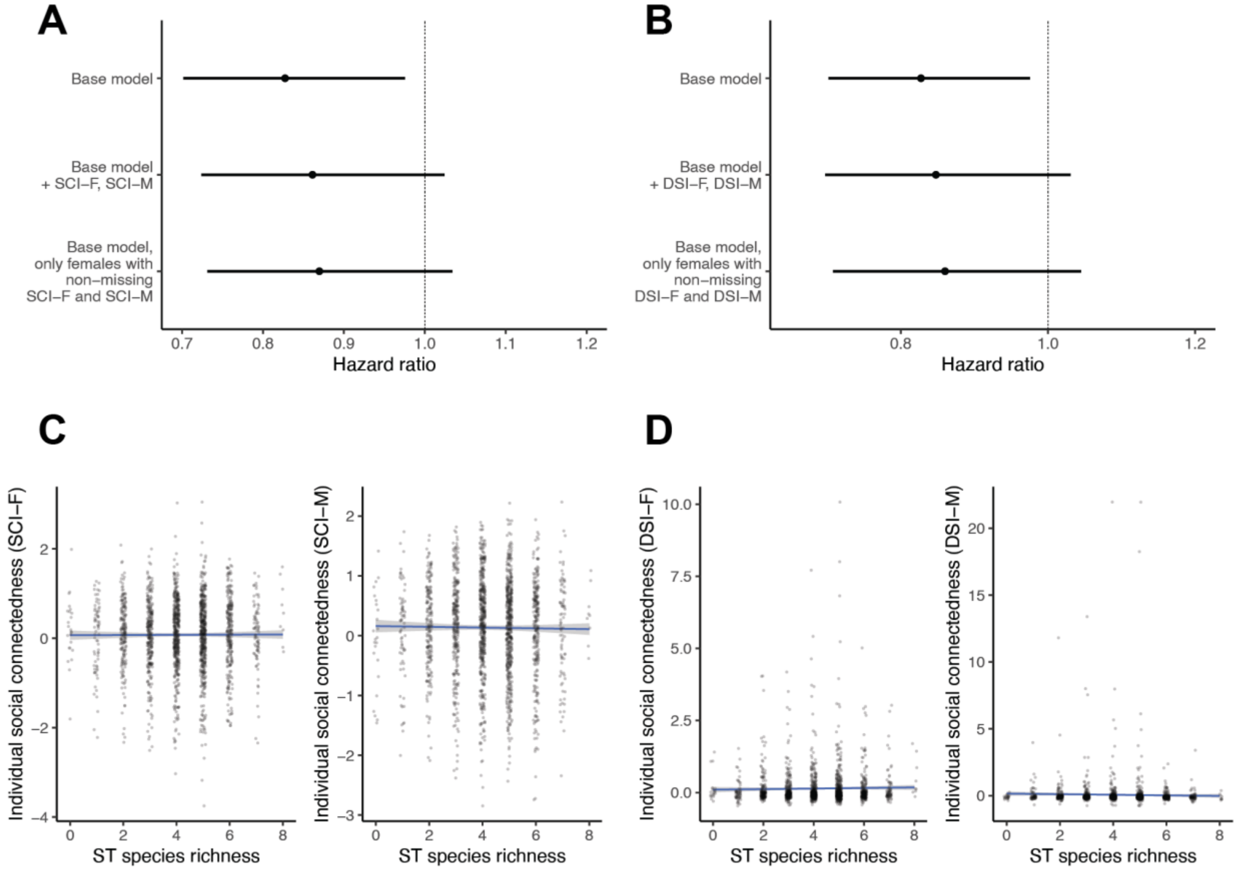


**Figure S21: Relationship between social connectedness, social bond strength, richness of socially transmitted species, and survival.**

**(a)** Adding $SCI_F$ and $SCI_M$ (observer effort-corrected grooming rates for females with all male or female group members, respectively) to the survival model does not substantially change the estimate of the relationship between the richness of ST species and female lifespan **(b)** Adding $DSI_F$ and $DSI_M$ (observer effort-corrected dyadic grooming rates for females with their top three female or male partners, respectively) to the survival model does not substantially change the estimate of the relationship between the richness of ST species and female lifespan. In both (b) and (c), adding the social variables causes the confidence intervals to overlap 1 due to reduced sample sizes of females with complete social integration/social bond data. **(c)** Socially transmitted species richness is not directly associated with $SCI_F$ or $SCI_M$ (linear mixed-effects model controlling for season and individual; $SCI_F$: ß=0.010, p=0.305; $SCI_M$: ß=0.0003, p=0.976). **(d)** We observe a slight positive relationship between ST richness and $DSI_F$, but not $DSI_M$ (DSI-F: ß=0.017, p=0.077; DSI-M: ß=0.003, p=0.885).

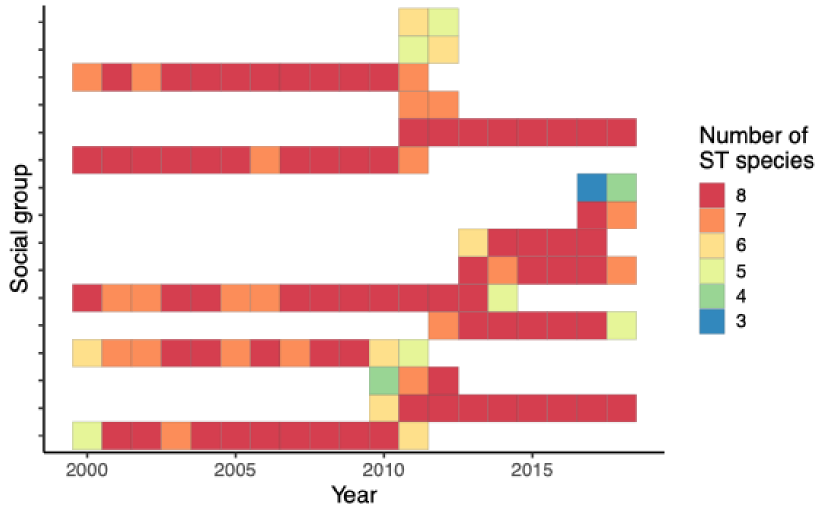


**Figure S22: Number of ST species detected per group in each year.**
ST species were considered present if they were detected in at least one member of the group at any point(s) in the year. The total number of circulating ST species was less than the theoretical maximum of 8 in one-third (33%) of group-years.

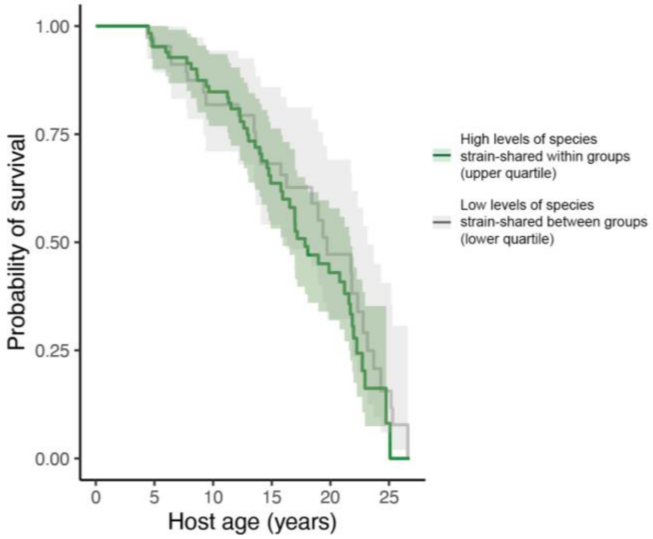


**Figure S23: Defining social transmission based on strain sharing within social groups does not identify a social transmission-linked signature of host survival.**
Female baboon survival is not predicted by the richness of species that are predominantly strain-shared within groups versus between groups, without additional information about source or timing of acquisition (hazard ratio=1.040, p=0.297).

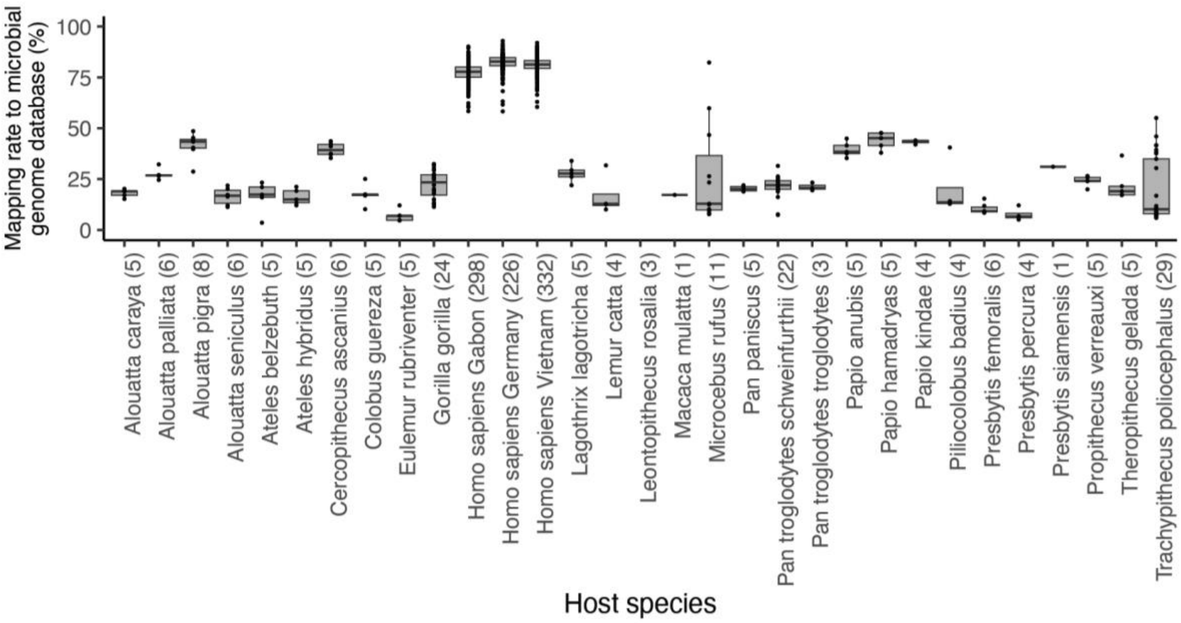


**Figure S24: Overview of primate comparative metagenomic dataset.**
Mapping rates of publicly available primate metagenomic datasets to our custom reference database of 4,712 microbial genomes. Sample size for each species is indicated in parenthesis on the x-axis labels.

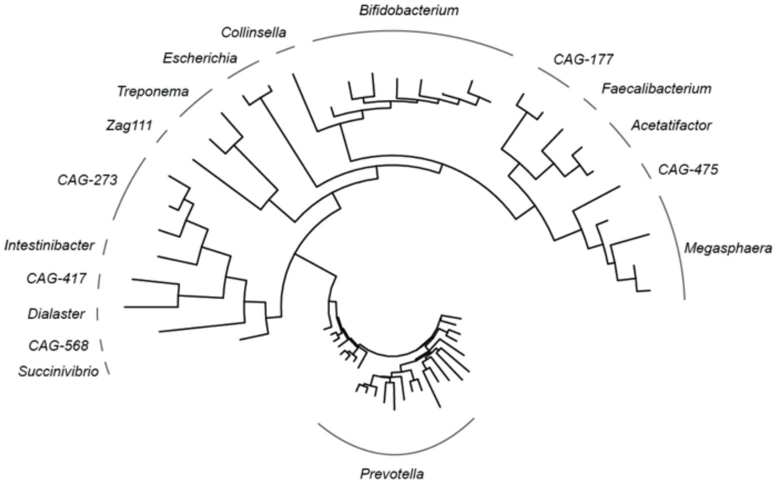


**Figure S25: Microbial species used in human co-diversification analysis.**
Phylogeny of all microbial species detected in gut metagenomes from humans living in Germany, Gabon, or Vietnam(6) that were from the same genus as one or more species implicated in a transmission event in the Amboseli baboons. Clades are labeled with genus names.
